# Unannotated translation products are widespread in model *E. coli*

**DOI:** 10.1101/2025.09.25.678689

**Authors:** Scott H. Saunders, Ayesha M. Ahmed, Manuel Razo-Mejia

## Abstract

Genomes contain orders of magnitude more open reading frames (ORFs) than known protein coding genes, and recent work suggests there may be unannotated proteins present in even the best studied organisms. To address this gap, we used a high throughput reverse genetic toolkit to construct precise C-terminal fusions of a reporter (and control) to >120,000 ORFs in model *E. coli*. We found hundreds of unannotated significant hits, and individually detected >50 novel polypeptides by western blot, including ORFs within tRNA loci. Many ORFs overlap annotated genes in the sense orientation, and we found these are likely chimeric polypeptides produced by ribosomal frameshifting. Using degron based knockdowns, we identified unannotated proteins that have putative fitness effects, and we found a novel small protein that displays phenotypes consistent with a role in the mRNA degradosome. The observation of a range of unannotated translation products should lead to better annotation and understanding of the bacterial domain of life and motivates the continued exploration of genomes broadly.

## Main Text

The accurate annotation of bacterial genome sequences remains an important challenge, which limits our understanding of both novel clinical and environmental isolates and long studied model organisms. Perhaps the most obvious genomic features worth annotating are protein coding genes, which are identifiable as open reading frames (ORFs) in DNA sequences. However, nearly every DNA sequence contains large numbers of possible ORFs, so how are true protein coding sequences differentiated? The simplest strategy has been to ignore short ORFs below a certain length cutoff. This is reasonable considering short ORFs appear frequently in random DNA, and there is a clear size difference between all possible ORFs and annotated genes in the model *Escherichia coli* genome (Fig. 1A)(*1*). However, recent work has repeatedly demonstrated that some short ORFs encode functional, unannotated proteins (*2–5*). Computational strategies to annotate protein coding genes have become more sophisticated, but most methods are limited by the few examples of small proteins identified in model organisms (*6, 7*).

**Figure 1.**
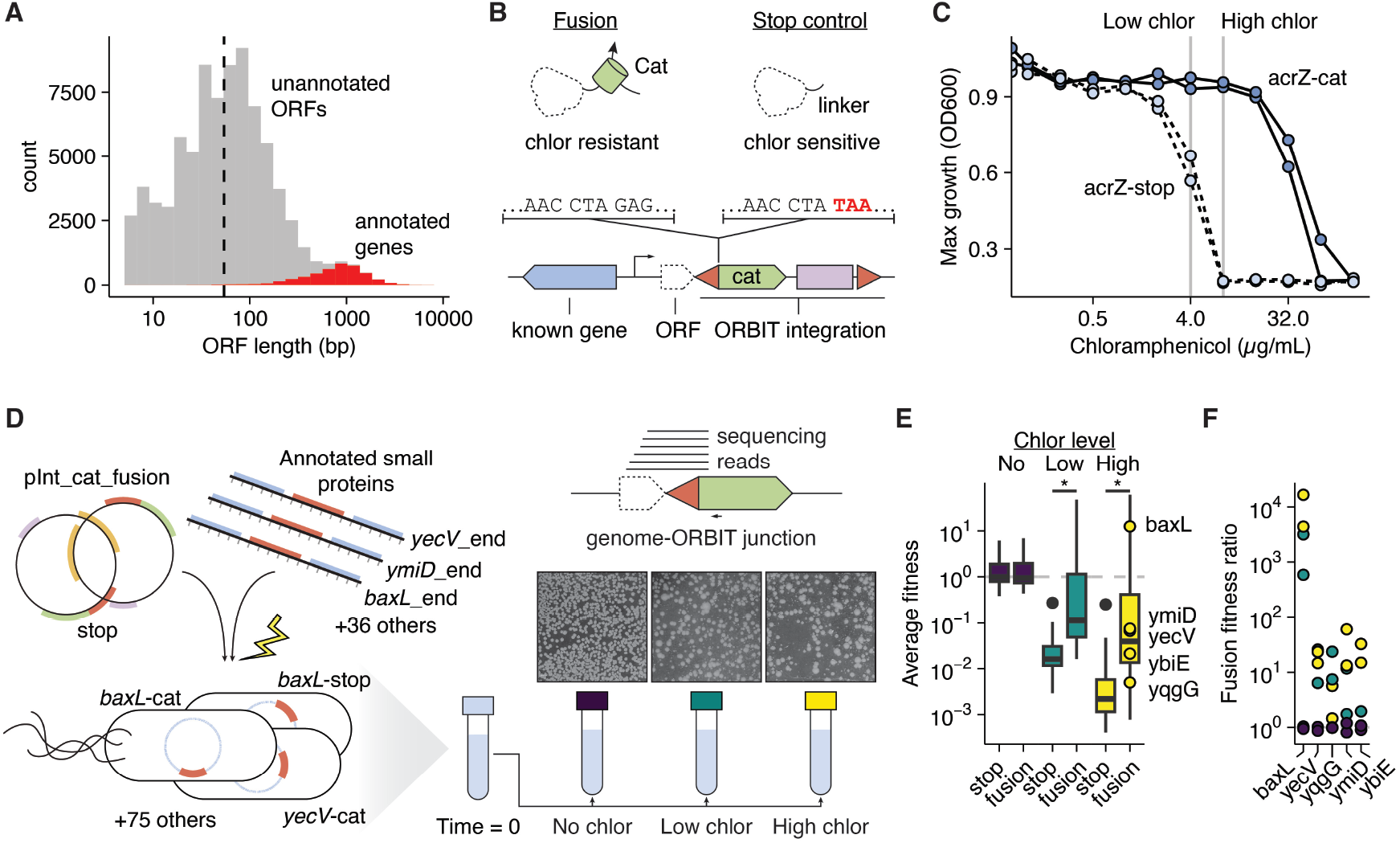
Establishing the CAT fusion approach. A) All ATG starting ORFs in the *E. coli* MG1655 genome are shown as length histogram. Annotated protein coding genes are highlighted in red. The dashed line represents the overall median length of 54 bp. B) The CAT fusion approach is shown, which confers chloramphenicol resistance upon ORF translation. The ORBIT genetic locus shows how the stop control is implemented in the linker region. C) The max growth achieved in 24 hrs (OD600) is shown for the *acrZ*-CAT fusion and strain and the stop control (n=2). Gray lines show the high (8 µg/mL) and low (4 µg/mL) chloramphenicol concentrations used throughout this work. D) A targeted mutant library was constructed with ORBIT using oligos targeting the C terminus of known small proteins and integrating plasmids with the CAT tag and stop control. Mutant libraries were selected on different concentrations of chloramphenicol, typically on agar, and genomic junctions were sequenced. E) Boxplots show the average fitness for the targeted fusions and stop controls in the different chlor concentrations. Asterisks denote significantly different mean values (t-test, unequal variance, p<0.05). F) Individual genes are highlighted to show FFR in different conditions.

Mass spectrometry-based proteomics and ribosome profiling (i.e. Ribo-seq), the sequencing of RNA associated with translating ribosomes, have revealed novel polypeptides leading to the “microprotein revolution,” which is now underway in diverse model systems in biology from plants, to cancer cells, viruses and microbes (*2, 5, 8*– *10*). These techniques are powerful, but mass spectrometry is poorly suited to confidently detect short polypeptide sequences. Ribo-Seq lacks definitive base pair resolution in bacteria, and it has inherent limitations around profiling translation associated loci. An orthogonal way to assess translation is with genetics. Random transposon systems with translational reporters have shown promise (*11, 12*), although genome coverage is limited, and polar effects can confound results.

To broadly survey the model *E. coli* genome for unannotated proteins, we chose to leverage precise high throughput genetics with our oligo-based method, Oligo Recombineering followed by Bxb-1 Integrase Targeting (ORBIT)(*13, 14*). We reasoned that high throughput reverse genetics could enable us to not only identify novel translated sequences, but also to perturb and manipulate them, gaining further insight and clues to function. ORBIT works by specifying modifications in DNA oligos with genomic homology arms, which direct the integration of a non-replicating plasmid that can contain various payloads. By using diverse pools of targeting oligos, we can rapidly design and construct mutant libraries with single nucleotide accuracy.

Here we generate a precise C-terminal ORF fusion library genome wide to create a new map of translation. We observed both annotated genes and unannotated ORFs and then made targeted ORBIT libraries of top hits to further test for translation and phenotype upon knockdown. Ultimately, we found compelling small protein hits at several loci, including core tRNA and rRNA genes, as well as ribosomal frameshift products overlapping annotated genes.

## Results

Our first goal was to develop a scalable sequencing-based assay to track fusions of a reporter to ORFs of interest. Because our precision genetics and tracking abilities scale efficiently (e.g. for libraries >10k mutants), we reasoned that this approach could enable us to ask if every possible ORF in the genome might be expressed as a protein.

### Pooled Cat fusion libraries detect known small proteins

To start, we chose chloramphenicol acetyltransferase (CAT) as a reporter, since it has been used for quantifying protein expression level (*15, 16*). The idea is simple: if a target gene is expressed as a protein, the CAT tag will also be translated and confer chloramphenicol resistance (Fig. 1B). We started with a model small protein, AcrZ, and used ORBIT to make a C-terminal fusion to CAT, along with a control containing a stop codon following the linker. The *acrZ*-CAT fusion strain grew well in high levels of chloramphenicol (> 16 µg/mL), while the stop control was completed inhibited at levels >4µg/mL, suggesting that our growth assay could be highly sensitive to protein expression at the right levels of chloramphenicol (Fig. 1C).

We next designed an ORBIT targeting oligo library to fuse CAT to a set of recently identified, low expression small proteins (*2*), which we thought would be highly representative of the type of new ORFs we hoped to find. We constructed both fusion and stop control pooled mutant libraries and assayed them at three different drug levels: 0 µg/mL (LB only), 4 µg/mL (low chlor), and 8 µg/mL chloramphenicol (high chlor) (Fig. 1D). By sequencing the CAT – genome junctions, we could precisely identify and quantify fusion and stop controls for 36 of the 38 small protein targets with high reproducibility (Fig. S1A). Initially these assays were performed in liquid medium, but we later found that growth on agar plates provided better selectivity and opportunity for individual strains to grow (Fig. S1C). It was readily apparent that we had fusions of different expression levels, since colony sizes varied dramatically in the presence chloramphenicol (Fig. 1D).

As expected, for nearly all loci, the CAT fusion increased in relative abundance compared to the stop control in the presence of chloramphenicol (Fig. 1E). This indicated that the CAT tag attached to each gene was translated into a functional protein and conferred a fitness advantage. We quantified this effect as a fusion fitness ratio (FFR), which compares the fitness of a fusion strain to the stop control. Therefore, no effect (i.e. FFR = 1) is observed without chloramphenicol, but FFR rises in the presence of the drug for strains that exhibit target gene translation. This assay proved to be strikingly sensitive, for example, *baxL* has a FFR of ∼10,000, meaning that the fusion was 10,000 times more fit relative to the stop control, and very low expression proteins were still identified with a FFR of ∼10 (e.g. yecV) (Fig. 1F, S1B)(*2*). Overall, 28 of 36 (78%) observed proteins yielded an average FFR greater than 2-fold in the high chloramphenicol condition (83% in low drug), while only 1 of 36 met this threshold without chloramphenicol (3%).

### Design and construction of a genome scale ORF fusion library

With a sensitive and scalable assay established, we decided to expand our approach to every possible ORF in the *E. coli* MG1655 genome. We first created a pipeline to find every ORF with the standard ATG start codon, as well as the common alternative start codons, GTG and TTG. This yielded 439,900 unique ORFs with different start sites, but only 137,800 unique stop codon sites, as most ORFs have 2-3 possible starts. The only limitation we set was ORFs had to be >= 15 bp, therefore generating polypeptides of at least 5 amino acids (including start methionine). Then for each ORF we computationally designed a C-terminal fusion targeting oligo of 120 nucleotides (nt), following our previously established design rules (*13*). These oligos evenly target ORFs across the genome, with an average spacing of ∼33 bp, including ORFs in every possible orientation and context (e.g. intergenic and genic) (Fig. 2A). We ordered these targeting oligos as subpools with different flanking primer sets, enabling us to separately process the annotated genes and unannotated sets of targeting oligos. Then we amplified and processed the oligos to obtain the desired products and transformed them into ORBIT cells in several rounds. Overall, we estimate we obtained >10 million recombinants, corresponding to at least 30x coverage of the 137,800 fusion and stop control constructs. Next we grew and pooled these mutant libraries and plated at desired densities on high, low, and no chloramphenicol giant agar plates (Fig. 2B). We performed this experiment and prepared sequencing libraries on two separate occasions for a total of 6 replicates.

**Figure 2.**
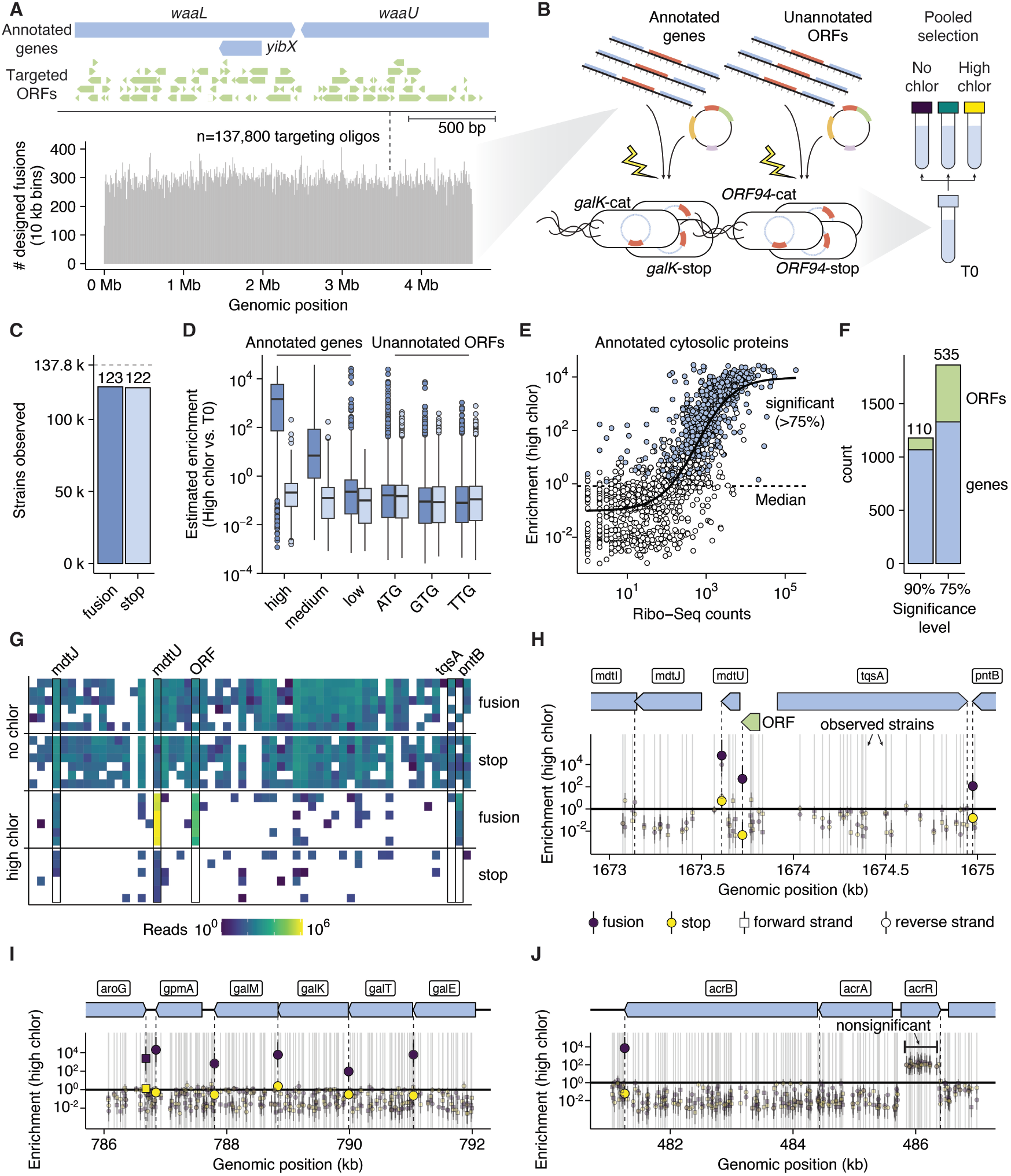
Constructing and measuring a genome wide fusion library. A) Genome wide targeting oligos are binned into 10 kb regions and counts are shown. Above, a single genomic region is shown as a demonstration, with each annotated gene (blue) and each ORF (green) targeted by an oligo. B) ORBIT libraries for annotated protein coding genes and ORFs were constructed separately and then pooled and selected on chloramphenicol. C) Across all sequencing reads, strains that were observed with at least 10 perfect reads are considered “observed” and counted in this plot. D) Enrichment values were inferred for each fusion and stop target. Targets were categorized based on the known expression level (annotated genes) or putative start codon (unannotated ORFs). E) Enrichment values for annotated cytosolic protein coding genes are compared with ribosome profiling data. A logistic model is fit to the data and shown as a black line. Significant hits are colored blue, while nonsignificant hits are colored white. F) Significant hits are shown for two different confidence levels 90% and 75%. G) For a single genomic locus containing *mdtU*, read counts for individual ORFs (columns) are shown across replicates and conditions (rows). H) The locus for data in figure G is shown as a diagram (top), and underneath are enrichment estimates for each observed strain containing a fusion or stop control in this genomic region. Gray lines represent observed strains, while points show enrichment values. Semi-transparent symbols are not significant hits, while larger solid symbols are significant. Squares denote forward strand targets and circles denote minus strand targets. I) Enrichment estimates for the *galK* locus is shown – no unannotated hits are called as significant. J) Enrichment estimates for the *acrR* locus is shown and the mutations within *acrR* are highlighted as nonsignificant.

From a total of ∼4 billion reads in the entire dataset, we observed 122,594 fusions and 121,952 stop controls with at least 10 reads perfectly matching to a designed genome-CAT junction, corresponding to an ∼88.7% success rate (Fig. 2C, S2A-E). Read counts across the replicates for each condition showed high correlations, indicating that the data was driven by reproducible effects (Fig. S2F-G). We observed 4,069 annotated gene fusions (95%), and many of those missing were highly essential genes. Interestingly, many essential genes did tolerate CAT fusions or the linker only (stop controls), for example *folA* and *rpoBC* (Fig. S4A-B). Ultimately, we believed that we had successfully made as many of the ORF-CAT fusions as reasonably possible.

### A genome wide map of translation identifies annotated genes and unannotated ORFs

To assess the significance of any unannotated ORF-CAT fusion, we calibrated our expectations from the set of annotated genes. On average, the annotated gene-CAT fusions (n ∼4,000) were enriched relative to their stop controls under chloramphenicol selection, while the unannotated ORFs did not show an overarching effect (n ∼ 118,000). To infer fitness in a rigorous manner, we adopted a hierarchical Bayesian approach, which was robust to zeros that are prevalent in the stringent high chloramphenicol condition (Fig. S3)(*17*). We categorized the annotated genes into high, medium, and low expression based on ribosome profiling data (*18*), and we observed a stepwise enrichment of the fusions with increasing expression level (Fig. 2D). Directly comparing Ribo-seq counts to chloramphenicol enrichment showed a clear sigmoidal relationship, indicating that our enrichment estimate likely corresponds well to protein copy number within a certain range (Fig. 2E). This is intuitive, since very low expression proteins would not produce sufficient CAT to impact intracellular chloramphenicol levels, and at extremely high expression, CAT levels may be sufficient to modify nearly all chloramphenicol, limiting resolution in this range.

To establish a significance cutoff, we used the enrichment and FFR values for the median annotated gene and then used conservative posterior predictive intervals to set probability cutoffs. Ultimately, we used 90% and 75% chance that a given strain was greater than the median gene levels, which respectively identified 1,068 and 1,329 annotated genes and 110 and 535 unannotated ORFs in the high chloramphenicol condition (Fig. 2F). At the more permissive 75% confidence level, our results were still highly stringent – identifying 69% of high expression, 45% of medium expression and 7% of low expression annotated genes. The low chloramphenicol condition identified similar numbers of hits, and there may be lower protein expression levels that this condition is better suited to reveal, but we focused on the high drug condition hits. We also found that simply examining the number of high chloramphenicol fusion replicates with reads was quite informative and used a hard cutoff of n > 3.

These data yield a genome wide map of translation that is highly informative. Figure 2G-H shows how reads correspond to enrichment scores and significant hits in a genomic context. This locus shows three significant hits – the annotated genes *mdtU* and *pntB*, and one unannotated hit, ORF2-16833. Generally, the genomic data correspond well to known operon structures, for example, our screen identified exactly 6 significant hits in the *galK* locus, which match perfectly to the known genes (Fig. 2I). Importantly, our stop control is also highly effective in several notable cases. For example, ORF fusions that inactivate the efflux repressor gene *acrR* confer strong chloramphenicol resistance by causing overexpression of the efflux pump *acrAB*, which natively acts on chloramphenicol. However, none of these ORFs were called as significant hits because the stop controls behaved the same as the fusions (Fig. 2J). There are other important reasons why CAT fusion fitness may report perfectly on protein expression level. The clearest example of this is outer membrane proteins, which must be exported from the cytosol, rendering CAT non-functional (Fig. S4C).

We observed diverse unannotated hits that occupy intergenic and genic regions, putatively use all three start codons (ATG, GTG, TTG), and occur throughout the genome. These ORFs are generally very short (median = 60 bp), although, some approach canonical lengths (i.e. >300 bp). One interesting set of hits occurs near important RNA genes that are not known to be translated. We found 10 unique tRNA loci that contain unannotated hits under the high chloramphenicol condition (Fig. S5). The 16s rRNA genes also show significant ORF hits, in the upstream region and a single hit near the terminus of the RNA gene (Fig. S6). Many other interesting unannotated hits and annotated genes are observed genome wide, so we have made this data readily accessible through a web-based genome browser interface (saunderslab.shinyapps.io/ecoli_ORF_CAT_fusions)(Fig. S7). Given the high significance cutoffs, this pipeline likely represents a lower bound for the true number of expressed unannotated ORFs under our conditions.

### Novel ORFs show additional evidence of translation

For any individual unannotated hit, further evidence is required to establish that an ORF is translated as the encoded protein. Using our significance calling pipeline and manual curation, we chose a set of 122 ORFs for further study. First, we wanted to validate our chloramphenicol enrichment results at high throughput, so we designed a fusion assay with a different reporter, mus musculus DHFR (mmDHFR), which confers resistance to the drug trimethoprim in *E. coli* (*19*). As with CAT and chloramphenicol, we found selective levels of trimethoprim with fusions and controls for the model small protein AcrZ (Fig. S8A-B). We created an ORBIT mutant library of our top hits fused with mmDHFR or the stop control and assayed them on trimethoprim plates (low, medium and high). Upon sequencing, we observed nearly all the designed strains (119 / 122 fusion = 97.5%) and saw that ORF fusions were on average statistically significantly enriched compared to their stop controls across each trimethoprim condition (Fig. 3C-F). Individually, 75% of the observed hits showed medium trimethoprim FFR > 1 (Fig. 3C), indicating that are our chloramphenicol assay results are relatively consistent with a different drug and reporter.

**Figure 3.**
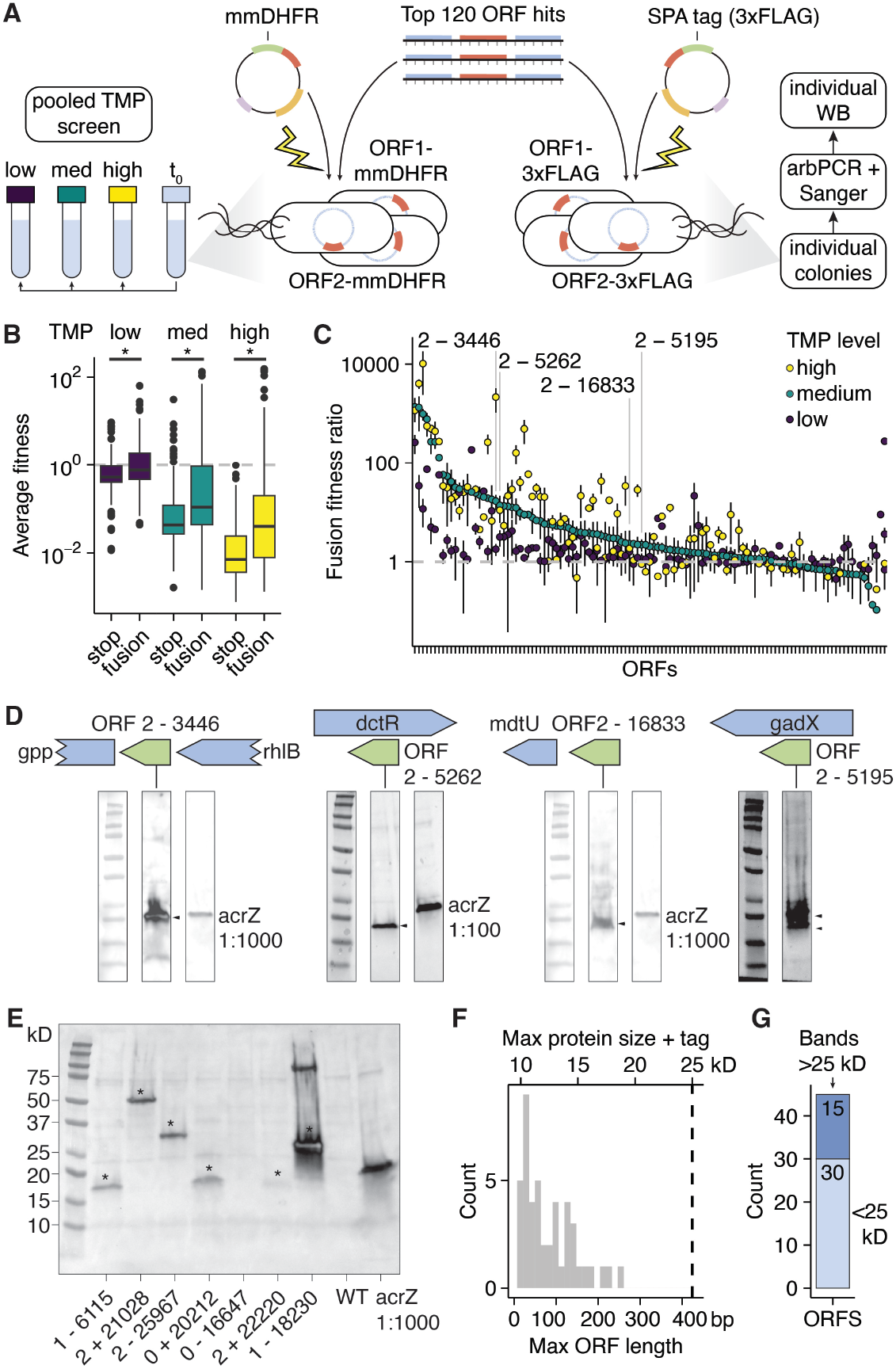
Further evidence of ORF translation. A) The diagram shows the approach to make targeted ORBIT mutant libraries with the reporter mmDHFR and selection with trimethoprim. The same targeting oligos were also used with an integrating plasmid carrying the SPA tag, which includes FLAG for western blotting. Western blotting was performed on individual strains that were identified following arbitrary PCR. B) Average fitness for each target is shown in the different trimethoprim (TMP) conditions. Asterisks denote significantly different mean values (t-test, unequal variance, p<0.05). D) The average (+/-standard deviation) FFR values are shown for all target ORFs. D) Western blots and loci diagrams are shown for four example unannotated hits. Arrows highlight the band specific to the tested ORF. E) An example western from the arbitrary PCR pipeline is shown. Asterisks denote bands not present in the WT control. F) The size distribution for all western blot observed ORFs is shown both in base pairs and the predicted molecular weight for the ORF with the SPA tag attached. G) The count of all western blot observed ORFs is shown, separately highlighting bands within the expected molecular weight and those that run higher than expected (>25 kD).

To further validate ORF translation, we used the set of 122 hit oligos to tag with the FLAG epitope for western blot (*2*). Construction was performed as a pooled library and individual colonies were arrayed and identified by arbitrary PCR and Sanger sequencing (*20*). By western blotting each strain with a FLAG antibody, we observed clear bands for >50 hits, strongly indicating that a novel polypeptide was produced from the genomic location and reading frame of our tag (Fig. S9). For example, ORF2-3446 is an intergenic (between *rhlB & gpp*) ATG starting ORF that is robustly expressed and runs at a very similar size to AcrZ (Fig. 2D). ORF2-5262 is a very short ORF (39 bp) encoded antisense to the gene *dctR*. ORF2-16833 is a clearly expressed ATG starting intergenic ORF running at the expected size. ORF2-5195 is a moderately expressed hit overlapping in the same direction (sense) as gadX, a lowly expressed gene. Interestingly, gadX appears as a doublet, which may indicate both possible start sites are used. These western blot results provided further evidence that most significant ORFs in our list of 122 genuinely produce novel polypeptides.

### Many sense intragenic hits are likely translational frameshifts of annotated genes

Upon western blotting many hits, we were surprised to see bands that were much higher than expected (Fig. 3E). Among the initial set of 45 ORFs we identified bands for, nearly all were shorter than 200 bp, so we established a conservative cutoff of 25 kD (ORF predicted size + tag) to decide if bands were significantly higher than expected (Fig. 3F). Fifteen hits were above this threshold, and each of them overlapped an annotated gene in the sense orientation (Fig. 3G), for example, ORF2+22177 overlaps *ilvC* and ORF2+25967 overlaps *deoD* (Fig. S10A-B). Looking more broadly at our set of high chloramphenicol significant hits, 245 of 535 ORFs were sense overlaps of an annotated gene, with a bias toward the +1 relative reading frame. These overlapping hits occurred throughout the annotated gene bodies and at varied expression levels (Fig. 4A). We tried to identify test cases for this class, for example, O antigen biosynthetic genes, *wbbI, wbbH* and *glf* form an interesting locus, where each has a strong unannotated ORF hit occurring in the +1 reading frame near the end of the annotated gene (Fig. 4B). Ribose-5-phosphate isomerase (*rpiA*) also contained a highly expressed ORF hit in the +1 frame near the end of the gene. We FLAG tagged the annotated gene and ORF for *wbbI, glf* and *rpiA* and observed clear bands much higher than expected based on the ORF size (Fig. 4C). Quantifying gel images revealed that the ORF products were 0.1 – 1% of the abundance of the annotated genes (Fig. 4D). We hypothesized that the high ORF bands could result from a ribosomal frameshift, where the upstream section of the annotated gene was fused to part of the ORF and the FLAG tag.

**Figure 4.**
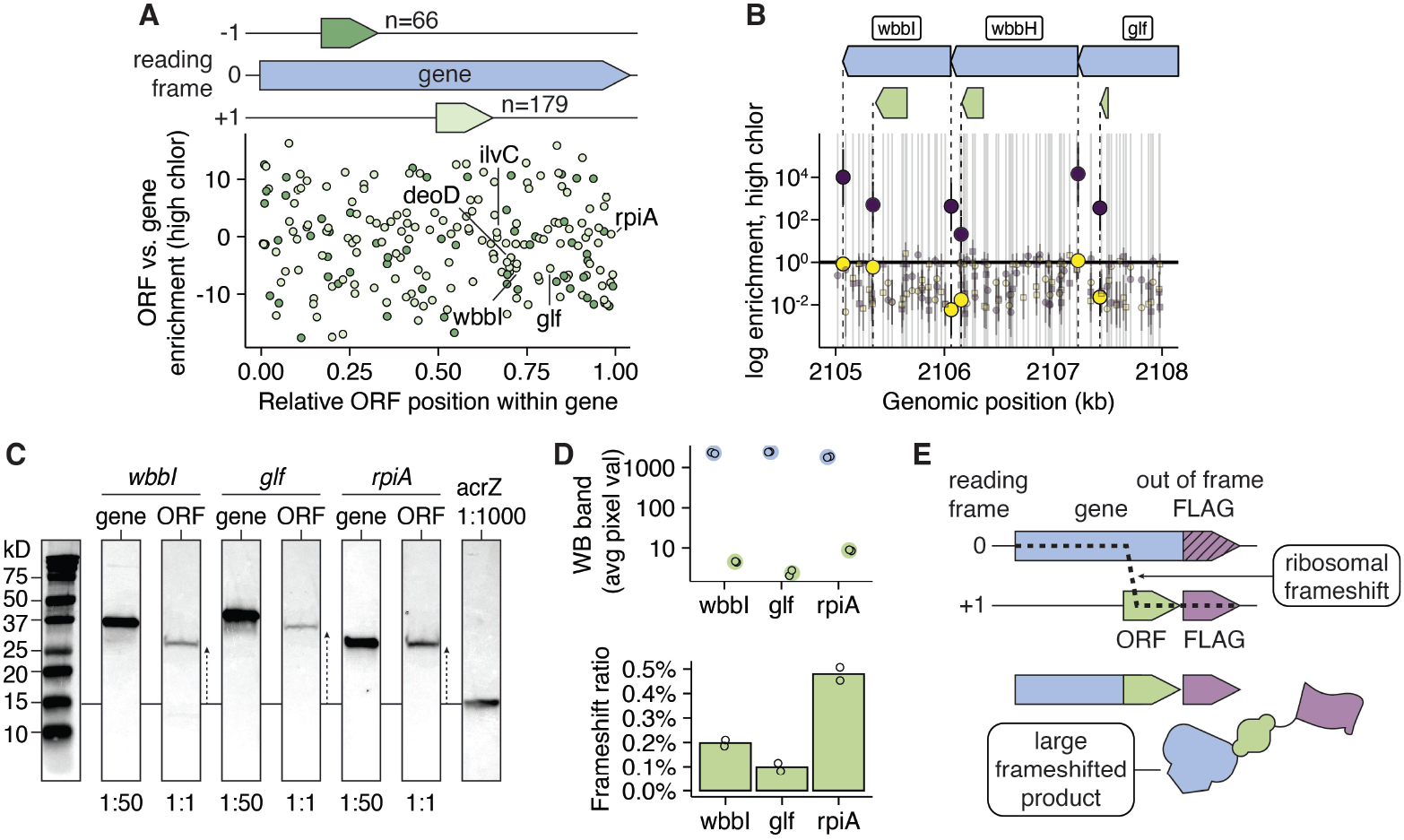
Frameshifting between annotated and ORF reading frames. A) High chloramphenicol significant hits (>75%) that sense overlap annotated genes are shown on the plot. The plot shows the ORF relative position within the annotated gene (standardized to length of 1) and it shows the relative enrichment level of the ORF relative to the annotated gene log_e_ scale. ORFs are colored by the relative reading frame, either + 1 or -1. B) An example locus containing wbbI is shown with chloramphenicol enrichment estimates. C) Three different example genes, wbbI, glf, and rpiA, and their sense overlapping ORFs were tagged and western blotted. Note annotated gene products were diluted 1:50, while the ORFs were not diluted to enable quantitative comparison. D) This schematic shows how a ribosomal frameshift can generate a larger than expected western blot product containing a FLAG tag.

To test if the observed bands originated from translation activity in the annotated gene reading frame (frame = 0), we precisely inactivated the start codons of *deoD* and *ilvC* by replacing them with a stop codon. These mutations were 508 and 982 bp upstream of the FLAG tag respectively (418 or 967 bp upstream of first possible ORF start respectively). These distant mutations eliminated the putative frameshift bands for *deoD* and *ilvC* (Fig. S10C). Conversely, replacing the start codon of the *ilvC* overlapping ORF with a stop codon did not eliminate the frameshift band, suggesting that the observed band does not initiate from the beginning of the ORF and the frameshift likely occurs downstream (Fig. S10D). Next, we wondered if the putative frameshifting rates were growth condition specific, so we grew *ilvC, deoD* and ORF tagged strains in LB and M9 conditions where *ilvC* and *deoD* are known to be important. The annotated genes both went up in abundance, but the frameshift products behaved differently. For ORF2+25967 (*deoD* overlap), the frameshift band disappeared, while for ORF2+22177 (*ilvC* overlap), the band increased in intensity more than the annotated gene (Fig. S10E-F). This suggests that the ORF translation products we observe may be condition dependent, and this process may be regulated through unknown mechanisms.

### Unannotated ORF perturbations show phenotypes

To test if any of our significant screen hits play functional roles in *E. coli*, we designed and implemented a degron based knockdown screen for phenotypes. First, we established that constitutive *ssrA* based degrons (recognized by *clpXP*) worked as expected (*21*). We added the degron tag (AANDENYALAA) to the ORBIT integrating plasmid and tagged three metabolic genes with obvious phenotypes, *galK* (galactose kinase), *metA* (homoserine O-succinyltransferase), and *leuD* (isopropylmalate isomerase subunit). Upon growth in selective media (i.e. with galactose, without methionine, without leucine), each degron strain showed profound growth rate defects, whereas the stop controls grew similarly to wildtype (Fig S11A).

Next we used our set of 122 ORF top hit oligos to make a degron and stop mutant library using ORBIT, and we screened the library in seven different liquid media conditions that represented diverse growth and stress conditions: LB (rich medium), M9 glucose (minimal medium), LB + H_2_O_2_ (oxidative stress), LB no shaking + NO_3_ (anaerobic respiratory growth), M9 glucose + NaCl (salt stress), LB + ciprofloxacin (DNA replication inhibitor), and LB + tetracycline (translation inhibitor) (Fig. 5A). We identified nearly all the designed strains and results were highly reproducible across the three replicates (Fig. S11C-E).

**Figure 5.**
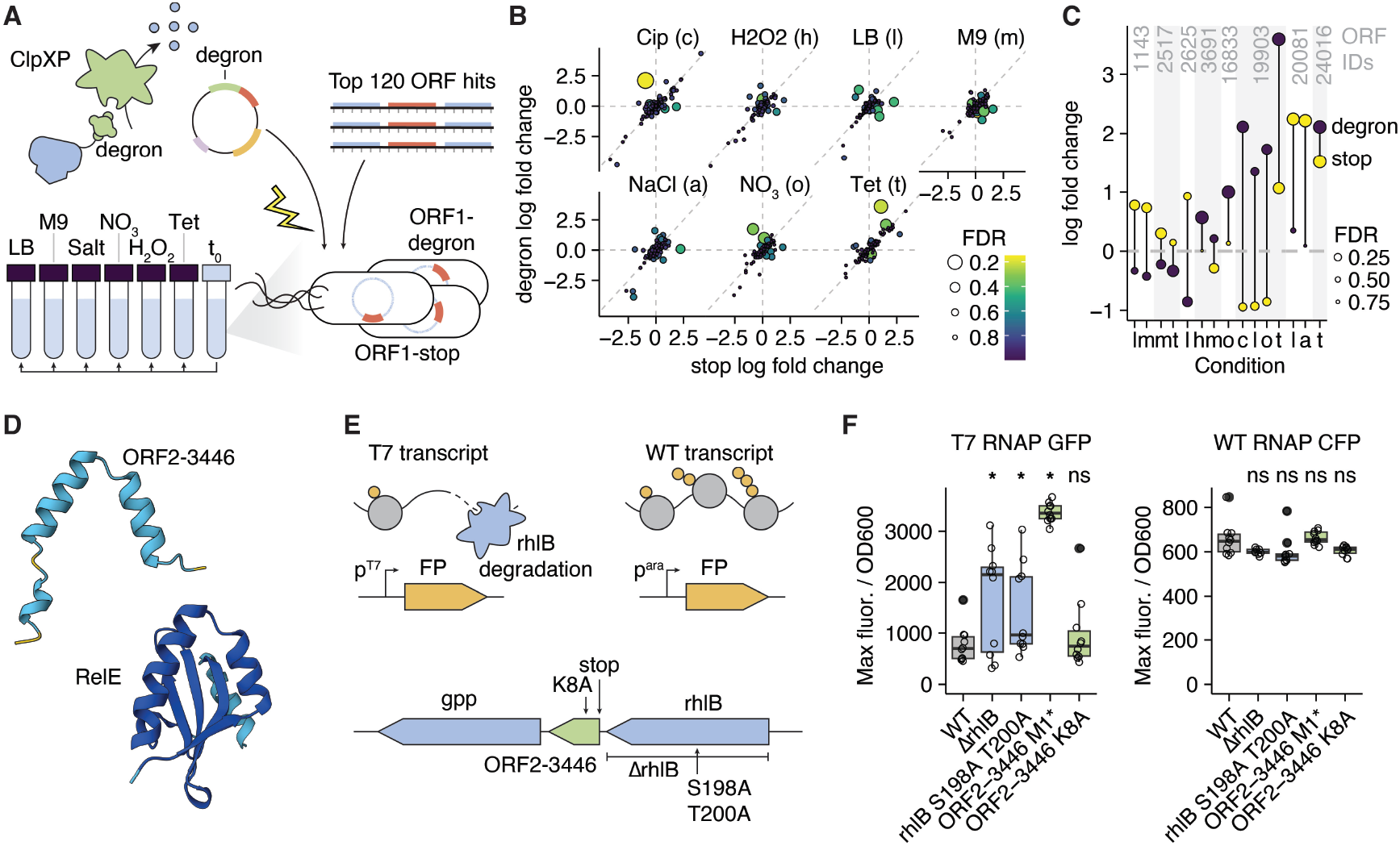
Degron screens and ORF2-3446 phenotype. A) This diagram shows how ORBIT was used to create a targeted degron library for screening phenotypes upon knockdown. B) Log fold changes are shown for degron and stop control strains in each of the seven tested conditions. The false discovery rate (FDR) of the degron vs. stop is shown as filled colors. C) Individual ORF – condition degron phenotypes are shown using one letter abbreviations for the different conditions. Here the FDR refers to the fusion log fold change or the stop log fold change only, not the contrast between the two. D) AlphaFold3 structures of ORF2-3446 and *E. coli* RelE are shown, colored by confidence (plDDT: dark blue = ‘very high’, light blue = ‘confident’, yellow = ‘low’, orange = ‘very low’). E) The ORF2-3446 locus is shown, including specific point mutations and a deletion of *rhlB* made with ORBIT (contains pInt_kanR integrated). The top portion shows the assay design, which utilizes fluorescent proteins expressed from a T7 RNA polymerase system or a standard arabinose inducible system. T7 transcripts are known to be relatively “ribosome free” due to the speed of transcription and are subject to degradation mediated by *rhlB*. F) Assay results are shown, as diagrammed in E. the maximum fluorescence per OD600 during a 24 hrs growth curve are shown, with the left plot containing the T7 results and the right plot containing a standard promoter / RNA polymerase (*araC* system). Asterisks denote significantly different mean values (t-test, unequal variance, p<0.05), while “ns” stands for “not significant.”

We observed significantly different effects for degron fusions and stops in each of the tested conditions (Fig. 5B), but ultimately a small number of effects (∼15) at a false discovery rate (FDR) < 0.5 (Fig. 5C). However, we realized that our stop control may still disrupt the function of small proteins, since the linker is still present. To test this, we went back and tested the degron on *acrZ* and showed that while the degron had the expected results (defect in chloramphenicol), the stop control was also defective (Fig. S11B). If we consider degron fusions with significant differences without comparing directly to stop controls, there are many more potential effects at a much more stringent FDR < 0.05 (>200 hits).

To investigate a novel hit in more detail, we selected ORF2-3446, which was one of our strongest hits across several preliminary experiments and shows modest effects in the degron screen. This ORF is completely intergenic, relatively large for a short ORF (123 bp), and appears to have a similar expression profile to the upstream gene, *rhlB*. Interestingly, blastP shows that ORF2-3446 has homology to the *E. coli* toxin RelE, which plays a role in cleaving mRNAs undergoing translation during stress (Fig. 5D)(*22, 23*). We hypothesized that a similar molecular activity might work with *rhlB*, which is an RNA helicase that is part of the mRNA degradosome and globally helps remodel and regulate mRNAs (*24, 25*). First, we introduced stop mutations into the ORF at the three possible start ATGs and confirmed that all three mutations eliminate the western blot band, indicating the upstream ATG is the true start codon. Next, we established an assay for mRNA degradation, based on known phenotypes for *rhlB*. GFP was expressed under control of T7 polymerase, which is known to generate ribosome free transcripts that are targeted by the mRNA degradosome (Fig. 5E)(*24*). We observed an effect for a deletion of *rhlB* and a point mutant inactivating RhlB catalysis. Inactivating the start codon of ORF2-3446 yielded a strong and consistent effect, whereas a site directed mutation in one of the conserved residues (from *relE*) did not (Fig. 5F). This clear phenotype suggests that this novel protein could play a role in mRNA degradation with RhlB, but further work is needed to characterize its molecular function.

## Discussion

Here we used high throughput ORBIT genetics to tag nearly all possible ORFs in the *E. coli* genome with a reporter and generated a genome wide map of translation that is methodologically orthogonal to existing proteomic and Ribo-Seq datasets. We confidently identified many annotated proteins that are known to be expressed under our culture conditions and established a conservative significance calling pipeline that identified over 500 unannotated hits. These data have been released in an accessible genome browser that will be a valuable unbiased resource for the community. For example, it is worth noting that many essential genes tolerate our translational fusions, and our results may offer useful estimates of expression for proteins that are difficult to detect by proteomics. Beyond our set of top identified hits, many more unannotated ORFs may be lowly translated considering our conservative significance threshold.

Closely examining annotated genes also reveals some of the shortcomings of our approach. First, ORBIT necessarily makes large modifications to endogenous loci, which limits our resolution of ORFs within highly essential genes and can cause unexpected secondary effects. We have rigorously controlled for these effects with the stop control, however, our method is likely to generate false negatives because of this. Further, the use of a translationally fused reporter may be suboptimal, since the CAT enzyme may function less efficiently when fused to certain proteins and complexes. One solution to these fusion issues may be to use inteins or other self-cleaving tags that could sever our reporter from the target proteins (*26*). One practical barrier to extending our high throughput genetic approach to other species is the cost and transformation efficiency required to target ORFs at genomic scales, although these barriers continue to lower with technological advances. Transposon mutagenesis may offer a way to perform initial screens in diverse organisms (*11, 12*), followed by more targeted ORBIT verifications. However, transposon mutagenesis is highly limited by its random nature and unavoidably “wastes” huge fractions of the mutant library on mutations within the same annotated genes.

We find many compelling unannotated hits that are worth studying individually. The ORFs overlapping the rRNA and tRNA genes stand out as particularly interesting and further work will be required to understand if they have true functions and what they might be. A priori, it seems plausible that these ORFs may play a role in regulation of their RNA components or the translational machinery.

It remains unclear what the true standard should be to establish an unannotated ORF as an annotated protein. These decisions are made even more complex by the growing understanding that many ORFs are evolutionarily young, and not necessarily broadly conserved (*27, 28*). We take the view that detected translation products should be generally annotated if they are detected, since ultimately biologists study extant strains that may incorporate any of these products into their physiology.

For a subset of 122 of our hits, we used targeted mutant libraries to further confirm ORF translation. However, we found that many of these are likely ribosomal frameshifts of annotated genes. Programmed ribosomal frameshifting is well documented, particularly in viral sequences, where molecular determinants like slippery codons and ribosome stalling mRNA structures are known (*29–32*). Interestingly, there are very few documented cases in *E. coli*, but they are notable. 1) *prfB* encodes release factor 2 (RF2), which is responsible for recognizing stop codons TGA and TAA. The gene has an in frame stop codon, which is suppressed in the absence of RF2 via a known frameshift, forming an important regulatory mechanism (*33, 34*). 2) *dnaX* encodes the tau and gamma subunits of DNA polymerase III, and gamma is produced by a programmed ribosomal frameshift (*35*). 3) *copA* encodes a copper transporter and a small chaperone protein CopA(Z) that is produced by a -1 frameshift (*36*). This work identifies many more putative endogenous frameshifting *E. coli* genes by western blot and CAT fusion screen. We generally observe lower frequency frameshifting than previously reported, but for high copy number genes we expect frameshifts to occur at 1-100 copies per cell, well within the range of other functional proteins. A variety of newer and older literature support our findings that ribosomal frameshifting could be wide spread at lower rates across the proteome (*29, 37, 38*). It is known the accuracy of translation is limited and error rates of frameshifting or misincorporation have been measured with individual reporters, but our dense genome wide screen provides a proxy measure of frameshifting across nearly every gene at multiple sites (*39*). Therefore, it remains unclear if our observed frameshifts are “programmed” or if the spurious frameshifting rate is just not even across sequences (*38*). Further mechanistic study will be required to deconvolve these ideas, but we suggest that evolution would be expected to act on the frameshifted proteins that we detect, and they could be functional and regulated. For example, a frameshift could cut off a domain of a protein, acting as a sort of isoform splicing in bacteria.

Overall, our work identifies widespread unannotated translation products in the best studied model organism and suggests other bacterial genomes may hold many more surprises. Beyond short ORFs, we believe that high throughput genetics offers a valuable way to re-examine our assumptions about genomes and to ask new questions in diverse bacteria.

## Acknowledgments

We would like to acknowledge Kimberly Reynolds and her lab for scientific support, feedback, access to laboratory equipment, and the pACYCDuet_WTDHFR_WTTS plasmid. We thank Boyuan Wang for feedback on protein biochemistry, and Siri Kothapalli and Grace Little for preliminary work. Thank you to the Lyda Hill Department of Bioinformatics and Green Center for Systems Biology for providing funding, as well as the computational resources of the BioHPC supercomputing facility.

## Author contributions

Conceptualization: SHS Formal analysis: SHS, MRM

Methodology: SHS, AMA, MRM Investigation: SHS, AMA

Visualization: SHS

Project administration: SHS

Supervision: SHS

Writing – original draft: SHS

Writing – review &editing: SHS, MRM, AMA

## Data and materials availability

All computational analysis is included as code notebooks in the github repository (github.com/saunders-lab/ecoli_short_ORFs). Plasmids will be available on Addgene or by request. Strains are available by request.

