## Supplemental materials for "Unannotated translation products are widespread in model *E. coli*"

### Supplementary Materials:

### Materials & Methods

#### *Cultures, cloning and strain verification*

Experiments were performed in *E. coli* K12 MG1655, referring to the genome sequence U00096.3. Strains were grown in LB liquid cultures in 14 ml round bottom polypropylene culture tubes in a shaking incubator (Innova 42) set to 37°C or 30°C (for temperature sensitive strains) and 225 RPM. Overnight cultures were typically started directly from –80°C frozen glycerol stocks. As needed, cultures were plated on LB + 1.5% agar (RPI #L42020). Both plates and liquid cultures contained antibiotics at the following concentrations: kanamycin sulfate 35 µg/ml, gentamicin sulfate 15 µg/ml, chloramphenicol 30 µg/ml (in 100% ethanol), ampicillin 100 µg/ml, carbenicillin 100 µg/ml. Carbenicillin was typically used in place of ampicillin due to its higher stability.

For chloramphenicol selections 4 µg/mL and 8 µg/mL were routinely used as the low and high concentrations in liquid and plates. Densities for plating were determined such that similar numbers were obtained across the conditions of different selectivity. For the genome wide CAT fusion experiment, giant agar plates were used (2x per condition replicate – Nunc Square Bioassay Dishes 245 x 245 x 25mm). The plates were incubated at 37°C for ~8 hrs and monitored closely to ensure colonies did not form a lawn. For trimethoprim selections, 0.0625 µg/mL, 0.125 µg/mL, and 0.25 µg/mL were used as the low, medium, and high concentrations in plates. For degron experiments, the growth conditions were LB, M9 glucose, M9 glucose + 2.5% NaCl, LB + 10mM $NaNO_3$ left in static incubator, LB + 500 µM $H_2O_2$, LB + ciprofloxacin 15.625 ng/mL, and LB + tetracycline 500 ng/mL. Fusion and stop mutant libraries were grown overnight in LB + kanamycin, then pooled and diluted 1:100 pooled in an LB culture grown to OD 1.0 (timepoint 0 – t0) and then diluted 1:100 into 3mL tubes of each condition of interest. Cultures were all grown in triplicate.

Plasmids were maintained in *E. coli* DH5a or pir+ hosts as appropriate. Integration plasmids were cloned from pInt_attP1_kanR (Addgene # 205294) and various sources, including IDT gBlocks, using the HiFi DNA assembly kit (NEB) to create pInt_attP1_cat_kanR, pInt_attP1_mmDHFR_kanR, and pInt_attP1_degron_kanR and their stop controls. The T7 reporter plasmid pACYC_T7RNAP_GFP_chlorR was cloned from pACYCDuet_WTDHFR_WTTS (Addgene #91226). Typically, helper plasmids were miniprepped (GeneJet) and integrating plasmids were midi prepped (Zymo).

Individually constructed strains with genomic modifications were checked by colony PCR with GoTagGreen G2 (Promega), and products were Sanger sequenced (Eurofins). Typically, these PCR products were "junctions" that were amplified using a primer on the genome and a primer on the ORBIT integrating plasmid. For ORBIT mutations (i.e. CAT, mmDHFR, degron & SPA fusions), 90 nt targeting oligos containing homology arms and the 38 bp attB site were typically ordered from IDT, although 120 nt oligos from Sigma Aldrich were also used. For point mutations introduced into ilvC, deoD, rhlB and ORF2-3446, 90 nt targeting oligos (IDT) were ordered containing only homology arms and desired mutations (no attB site) and transformed into ORBIT competent cells carrying the V1 helper plasmid, which can create SNPs due to the inclusion of MutL. Colonies were screened for mutations using a primer where the 5 prime end landed directly on the mutation. Positive colonies were restreaked and locus spanning colony PCRs were performed and sequenced to verify mutations. Phenotypic assays were typically performed in a Biotek multiplate reader (placed in 37°C warm room), monitoring OD600 and sometimes GFP fluorescence.

***ORBIT***

ORBIT competent cells were prepared following previously published protocols (*13*). Briefly, strains with the helper plasmid (pHelper_TS_V2_ampR Addgene #214467 or pHelper_Ec1_V1_gentR Addgene #205291) were grown overnight with antibiotic (3 ml cultures). In the morning, cultures were diluted 1:1000 into larger volumes, typically 500 ml, with LB + antibiotic. Cultures were grown ~3.5 h to OD ~0.3 and cultures were induced for oligo recombineering with 1 mM m-toluic acid (Sigma Aldrich #T36609) (1 M stock in 100% ethanol). After a 30 min induction continuing in the shaking incubator, flasks were put on ice and swirled periodically for ~15 min. Cultures were distributed into 50 ml falcon tubes and centrifuged at 5000 rcf for 10 min in a pre-chilled centrifuge. Tubes were transported on ice to a cold room, supernatant was removed, and pellets were thoroughly resuspended in ice cold milliQ water with a serological pipettor (50 ml and then 10 ml). Tubes were inverted 5 times, then transported on ice and centrifuged again. This process was repeated two more times with 10% glycerol washes. The final wash supernatant was poured off and cells were resuspended in residual buffer typically yielded 50–100× concentrated cells in 10% glycerol. These cells were aliquoted into microfuge tubes placed open faced on ice (50 µl aliquots). Each aliquot was then chilled for >5 min on dry ice before placing in freezer boxes and placed at –80°C. Competent cells typically retained high efficiency for several months.

To transform cells, frozen induced competent cell aliquots were taken from the –80°C box and thawed on ice for ~10 min or until completely thawed. Then targeting oligo (2 µl of 25 µM stock–1 µM final) and integrating plasmid (1 µl of 100 ng/µl) were added to each aliquot and aliquots were mixed 3 times with a 200 µl pipette before transferring to ice cold 0.1 cm electroporation cuvettes (Fisher #FB101). These conditions were standard for individually constructed strains and mutant libraries constructed with IDT oPools (Figure 1, 3 & 5), but Twist oligo pool mutant libraries differed (Figure 2) – see below. Oligonucleotides were resuspended in molecular grade water and used directly. Cuvettes were tapped and inspected to ensure no bubbles or gaps were present and electroporated with standard E. coli settings (1.8 kV, Bio-Rad MicroPulser). With a 1000 µl pipette, 1 ml of recovery media (LB + 0.1% l-arabinose) was gently added to the cells and cells were transferred to the 3 ml recovery culture tubes. Cultures typically recovered at 37°C shaking for 1 hour before plating.

***Ultra high throughput Twist oligo pools***

All ORFs in the genome were found using a custom R script (see github repository) and several loci were cross referenced with the Benchling ORF finder to confirm the accuracy. ORFs were numbered according to the reading frame and genomic strand. Therefore 2-27 would be reading frame = 2, strand = "-", number = 27, so all three pieces of information are required to uniquely specify a target ORF. Unique ORFs were considered only based on their stop codon position, so each ORF can have multiple possible start codons with ATG, GTG or TTG.

For each target ORF, an R script was used to design 120 nt C terminal targeting oligos that bind the lagging strand template for each. The resulting targeting oligos containing upstream and downstream homology for each target ORF (and the attB site). The targeting oligo sequences were separated into 14 subpools categorized by annotated status, attB direction (complementary attB sites were separated into different subpools to avoid PCR issues), expression level (annotated genes) and putative start site (unannotated ORFs). Next, we added orthogonal 20 nt primer binding sites and internal MluCI and BtsI-V2 recognition sequences. These sites were designed such that the targeting oligo sequences could be amplified with PCR, but then the primer sites could be entirely cleaved off. Oligos were ordered in a large pool from Twist Biosciences.

Upon receiving the oligo pool, it was resuspended according to the manufacturer's guidelines and subpools were independently amplified using their specific orthogonal primer sets. PCR was performed in two stages and different cycle numbers were tested to use as few cycles as possible and limit unwanted products. Products were assessed on 2% agarose gels. PCR1 used 1 µl of the original oligo pool diluted 1:20 with 20 cycles (Q5 hotstart polymerase, 98°C – 15 s, 64°C – 20 s, 72°C – 20 s). PCR2 was performed with 10 ng of purified PCR1 product for 7 cycles (same parameters as PCR1). Note that the forward primers had two phosphorothioate bonds at the 5′ end and the reverse primers had 5′ phosphate groups. These modifications promoted the degradation of the reverse (i.e. bottom) strand by lambda exonuclease. This digestion was performed on 1–3 µg of the PCR2 product with 1 µl of exonuclease (NEB #M0262) at 37°C for 30 min before heat inactivation. Digestion products (presumably single stranded DNA) were purified with the oligo clean and concentrate kit (Zymo #7010).

Next guide primers were annealed to the single stranded DNA (targeting oligo with primer overhangs), making the restriction sites duplex DNA and therefore suitable substrates for endonucleases. Annealing was performed with 4 µM (final) of each guide oligo and 0.5–2 µg of template DNA in 1x Cutsmart buffer (NEB) using a slow temperature gradient (95–60°C at 0.1°C/s, 60°C for 3 min, 60–50°C at 0.1°C/s, 50°C for 3 min, 50–37°C at 0.1°C/s, 37°C for 3 min). Then 5 units (1 µl) of each restriction enzyme (MluCI and BtsI-V2) were added to the 50 µl reaction and left at 37°C for 2-3 hours. Products were purified using the oligo clean and concentrate kit and loaded on 10% TBE–urea gels (BioRad #4566033) with sample buffer (Thomas Scientific #C995V39) and run at 200 V for ~30 min. Gels were stained with 1× Sybr gold (ThermoFisher #S11494) for 20 min and then imaged. For size comparison, 120 nt and 90 nt targeting oligos were run alongside digestion products. This process typically yielded incomplete products, but we proceeded with the heterogeneous reaction products, which worked acceptably well.

These processed targeting oligos were used for a standard ORBIT experiment with 50 µl induced electrocompetent cell aliquots. Approximately 100 ng of integrating plasmid and 2-4 µl of targeting oligo product was added to each transformation. Typically yields of these amplified and processed oligos were much less than individually ordered commercially synthesized oligos, so 1 µM final oligo concentration was rarely achieved. Following transformation, recovery cultures were concentrated and spread plated on multiple kanamycin plates to obtain large numbers of colonies. Colonies were resuspended in LB and pooled glycerol stocks were made.

#### *Sequencing library preparation and analysis*

Library preparation for ORBIT mutant pools was performed in a manner analogous to Tn-Seq (*40*, *41*). Genomic DNA (gDNA) was extracted from mutant library cell pellets using a gDNA miniprep kit (Zymo). DNA was fragmented and adapters were ligated using the NEB Ultra II FS Library Prep Kit for Illumina (#E6177). Fragmentation was performed for 10 min at 37°C and magnetic bead size selection was performed for fragments sized 275 to 475 bp. PCR1 was performed with 500 ng of adapter ligated DNA using a primer that binds the integrating plasmid and a primer (with overhang) that binds the adapter using Q5 hotstart polymerase (98°C – 15 s, 62°C – 20 s, 72°C – 30 s, 15 cycles). To avoid bottlenecking of diverse genome wide CAT fusion libraries, 4 replicate fragmentation / adapter ligation reactions were performed for each biological replicate condition. Illumina compatible overhangs were added with PCR2 using 1 µl of the unpurified PCR1 product (98°C – 15 s, 63°C – 20 s, 72°C – 30 s, 15 cycles). Products were cleaned with magnetic beads and served as the final sequencing library. Libraries were quantified with Qubit fluorometry (Thermofisher # Q32851) and typically pooled 1:1. For oPool mutant libraries (Figures 1, 3 & 5), sequencing libraries were run on an Illumina MiSeq instrument with a Nano V2 kit (300 cycles total, typically 200 cycles Read 1). For larger Twist mutant libraries, pooled sequencing libraries were sent to Azenta and run on NovaSeq instruments.

First, demultiplexed sequencing reads were quality filtered using default settings and fastp (*42*). Then ORBIT mutations were identified by treating them as adapters in Cutadapt (*43*). For fusions and stop controls, they were treated as separate adapters and Cutadapt categorized reads into separate fusion and stop control files. Following trimming of the ORBIT sequences, putative genomic sequences were mapped to the *E. coli* genome. Then genomic positions were matched between mapped reads and designed constructs to count the number of reads observed for each targeted strain. For MiSeq experiments, sequencing reads were processed on a standard desktop computer. For NovaSeq experiments, sequencing reads were processed on the BioHPC on a node with 128 GB of RAM and 32 logic cores. The github repository contains the sequencing read processing scripts.

To obtain fitness values for each mutant, relative abundances within condition replicates were calculated, and then each condition was compared with t0. Fusion fitness ratios were then calculated by further taking the ratio of the fitness for each fusion and paired stop control.

For the genome wide CAT fusion experiment (Figure 2), a hierarchical Bayesian framework was used to estimate enrichment across the 6 replicates. Briefly, a normal distribution was used to estimate the log frequence ratio mean and standard deviation for each target strain. The noise in read counts for each strain was estimated as a log normal distribution. Hyperparameter estimation was performed using the NumPyro package with stochastic variational inference and "Trace_ELBO." Posterior predictive intervals were calculated using the following relationships:

sample_log_freq ~ Normal(log_freq_ratio_loc, log_freq_ratio_scale)

sample_sigma ~logNormal(sigma_loc, sigma_scale)

sample_error ~ Normal(0, sample_sigma)

log_freq_ratio_ind ~ sample_log_freq + sample_error

Using the estimated log frequency ratios (i.e. estimated enrichment), significance cutoffs were set using the median annotated gene values for both fusion enrichment and fusion – stop enrichment (i.e. FFR). For each metric, the fraction of the posterior predictive interval above the cutoff was used as the confidence score (75% or 90%). To be called as significant, target fusions also had to be observed in at least three separate high chloramphenicol replicates.

For the degron experiments, edgeR was used to calculate fusion and stop specific log fold changes and false discovery rates, as well as a contrast of Fusion – Stops (equivalent to FFR) using the glmQLFTest (*44*).

#### *Western blotting and protein analysis*

We constructed a pooled mutant library tagging the set of 122 top hit ORFs with the sequential peptide affinity tag (SPA, encoded on ORBIT integrating plasmid), which includes both FLAG and calmodulin binding protein, for a total of 255 bp total. Then individual colonies were picked into 96 well plates and an arbitrary colony PCR protocol was performed. Briefly, GoTaq Green G2 (Promega) was used for colony touchdown PCR with two semi random forward primers (with an added overhang site) and an ORBIT specific reverse primer. A second enrichment PCR was performed with primers specific to the ORBIT modification and the overhang added in PCR1 (*20*).

Arbitrary PCR products were sequenced by Sanger (from ORBIT modified side) and data were aligned to the SPA tag template and genomic sequence was matched to the designed targets. Only perfect matches were included. Colonies were inoculated overnight in LB + Kanamycin and then diluted 1:1000 into LB before pelleting at OD ~1.0. Pellets were stored at -80°C if not used fresh. Pellets were resuspended in 100 µL of Laemmli Buffer + 5% fresh Betamercaptoethanol and boiled in microfuge tubes for 20 min. Any kD mini protean TGX 10 well gels were used from Bio-Rad (#4569033). Gels were run in 1x SDS PAGE buffer with Precision protein plus ladder (Bio-Rad). Typically, 25 µL of sample was loaded into each well. Gels were run for 30-40 min at 200V and then transferred to Nitrocellulose membranes. Membranes were dried for 15 min, then blocked with 10 mL TBST + 3% milk on an orbital shaker overnight at 4°C. An HRP conjugated primary anti-FLAG antibody (1:100) (Abcam #ab49763) was used to probe membranes by incubating 1 hour at room temperature on the orbital shaker. After washing 3x with TBST (15 min each at room temperature), chemiluminescent reagent was added (SuperSignal West Atto – Thermo Scientific #A38554). Membranes were placed in a sheet protector and imaged on a Bio-Rad ChemiDoc sytem. Typically, a strain with the *acrZ*-SPA tag was used as a control, diluted 1:100 or 1:1000 and an untagged wildtype strain was included to assess background bands.

The AlphaFold3 server was used to predict structures for ORF2-3446 and RelE (*45*).

**Supplemental figures**

Figure S1 Pilot CAT fusion details

Figure S2 Genome wide CAT fusion details

Figure S3 Enrichment inference and significance pipeline

Figure S4 Essential gene and localization constraints

Figure S5 Unannotated hits overlapping tRNA loci

Figures S6 Unannotated hits overlapping 16S rRNA loci

Figures S7 All other significant (>90% confidence) high chloramphenicol unannotated hits

Figures S8 mmDHFR fusion assay details

Figures S9 Western blotted ORFs

Figures S10 Ribosomal frameshift details

Figures S11 Degron assay details

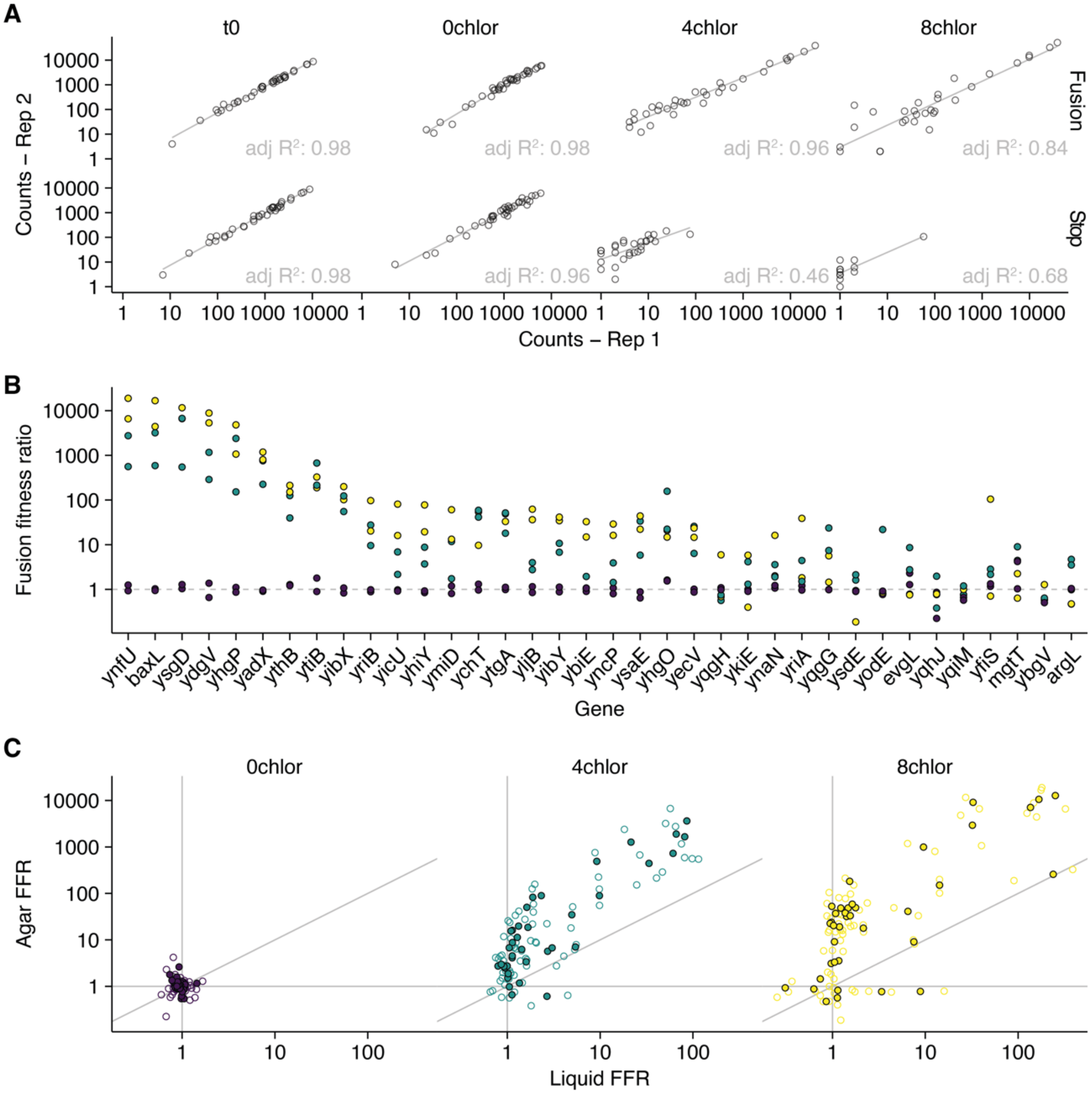


Fig. S1 Pilot CAT fusion details. A) Read counts are shown for replicates 1 and 2 for each sequenced library, broken down into fusion and stop control strains. Lines are linear models and adjusted $R^2$ values are shown in the bottom right corner of each plot. B) FFR for all genes are shown, the full version of main Fig. 1F. C) The entire assay was performed with selection in liquid media and on agar plates. The FFR for each target is shown for both experiments. Solid points show mean values, while open circles show individual replicates. The diagonal gray line is slope = 1, intercept 0, therefore points above this line show higher FFR in agar than liquid.

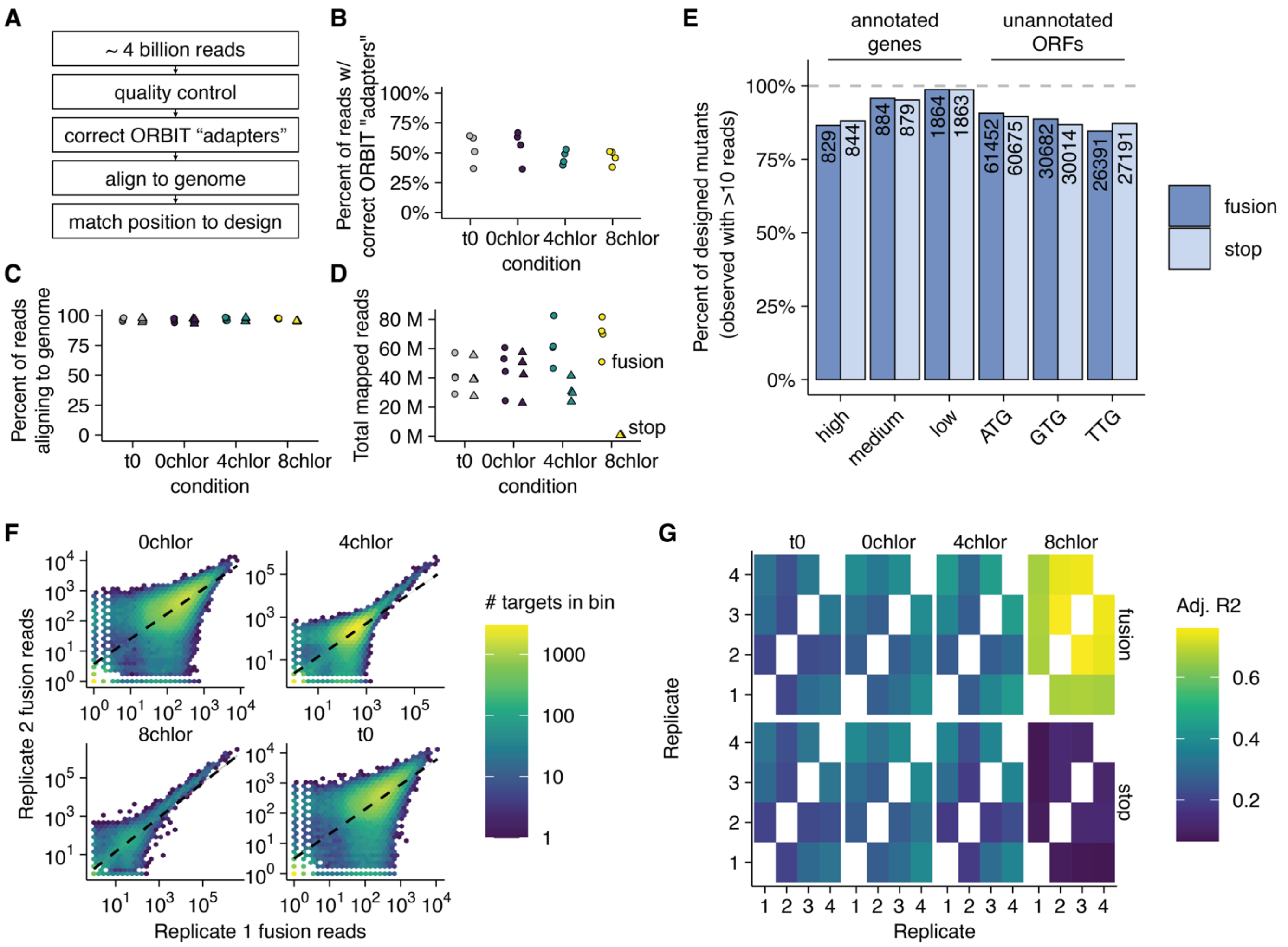


Fig. S2 Genome wide CAT fusion details. A) Sequencing analysis pipeline. B) The fraction of each sequencing library with the identified ORBIT sequence, treated as "adapters". Individual points are replicates. C) Of the trimmed reads, this plot shows the percent of reads that mapped to the E. coli MG1655 genome. Circles are fusions and triangles are stop controls. D) The total mapped reads for each library are shown, where M is millions of reads. E) The fraction of each subpool that was observed with at least 10 reads across the dataset. The individual number of strains observed within each group is shown as the numeric label at the top of each bar. F) Example correlations for fusions only in replicates 1 and 2. Read counts are the axes and the color is the number of targets in each bin. Black dashed lines are linear models. G) The adjusted $R^2$ values for all replicates / conditions are shown.

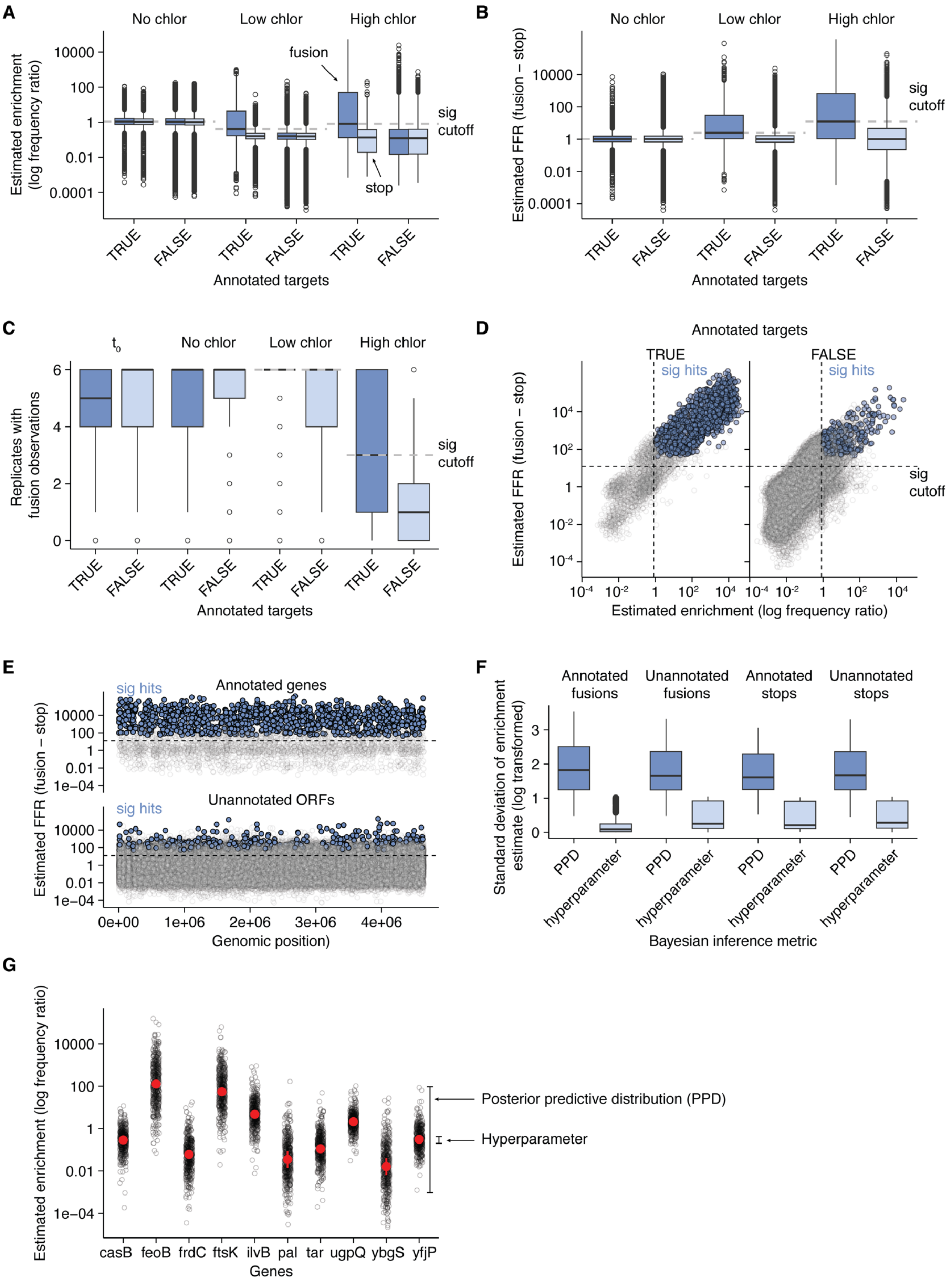
A
No chlor
Low chlor
High chlor
fusion
stop
sig cutoff
Estimated enrichment (log frequency ratio)
TRUE
FALSE
Annotated targets
B
Estimated FFR (fusion – stop)
C
t0
Replicates with fusion observations
D
Annotated targets
sig hits
Estimated enrichment (log frequency ratio)
E
Annotated genes
Unannotated ORFs
Genomic position
F
Annotated fusions
Unannotated fusions
Annotated stops
Unannotated stops
Standard deviation of enrichment estimate (log transformed)
PPD
hyperparameter
Bayesian inference metric
G
casB
feoB
frdC
ftsK
ilvB
pal
tar
ugpQ
ybgS
yfjP
Genes
Posterior predictive distribution (PPD)
Hyperparameter

Fig. S3 Enrichment inference and significance pipeline. A) Estimates of enrichment (log frequency ratio) are shown for fusions and stops, annotated genes and unannotated ORFs at different chloramphenicol concentrations. Gray lines are the significance cutoffs used, which correspond to the median values for annotated fusions. B) Estimates of fusion fitness ratio (fusion – stop enrichment) for annotated genes and unannotated ORFs. Gray lines are the significance cutoffs used, which correspond to the median values for annotated genes. C) The number of experimental replicates in which a fusion strain was observed (max of 6), shown for annotated genes and unannotated ORFs across chloramphenicol conditions and time zero. Gray lines show significance cutoffs (inclusive), which correspond to median values for annotated genes. D) The correlation between the two primary metrics, enrichment and FFR, showing significance cutoffs and identified hits for annotated genes and unannotated ORFs for the high chloramphenicol growth condition, >75% confidence level. Solid blue points are significant hits, while semitransparent gray points are not significant hits. E) The genomic distribution of significant hits is shown with FFR estimates. Solid blue points are significant hits, while semitransparent gray points are not significant hits. Dashed lines show the FFR significance cutoff. F) The Bayesian parameter estimation could be presented in two different ways – with only the hyperparameter or with a posterior predictive distribution. This plot shows the standard deviations of the two estimates for annotated genes and unannotated ORFs, fusions and stops. G) Example genes showing samples from the posterior predictive distribution (PPD) as black points, and the hyperparameter estimate with standard deviation shown as red points and lines.

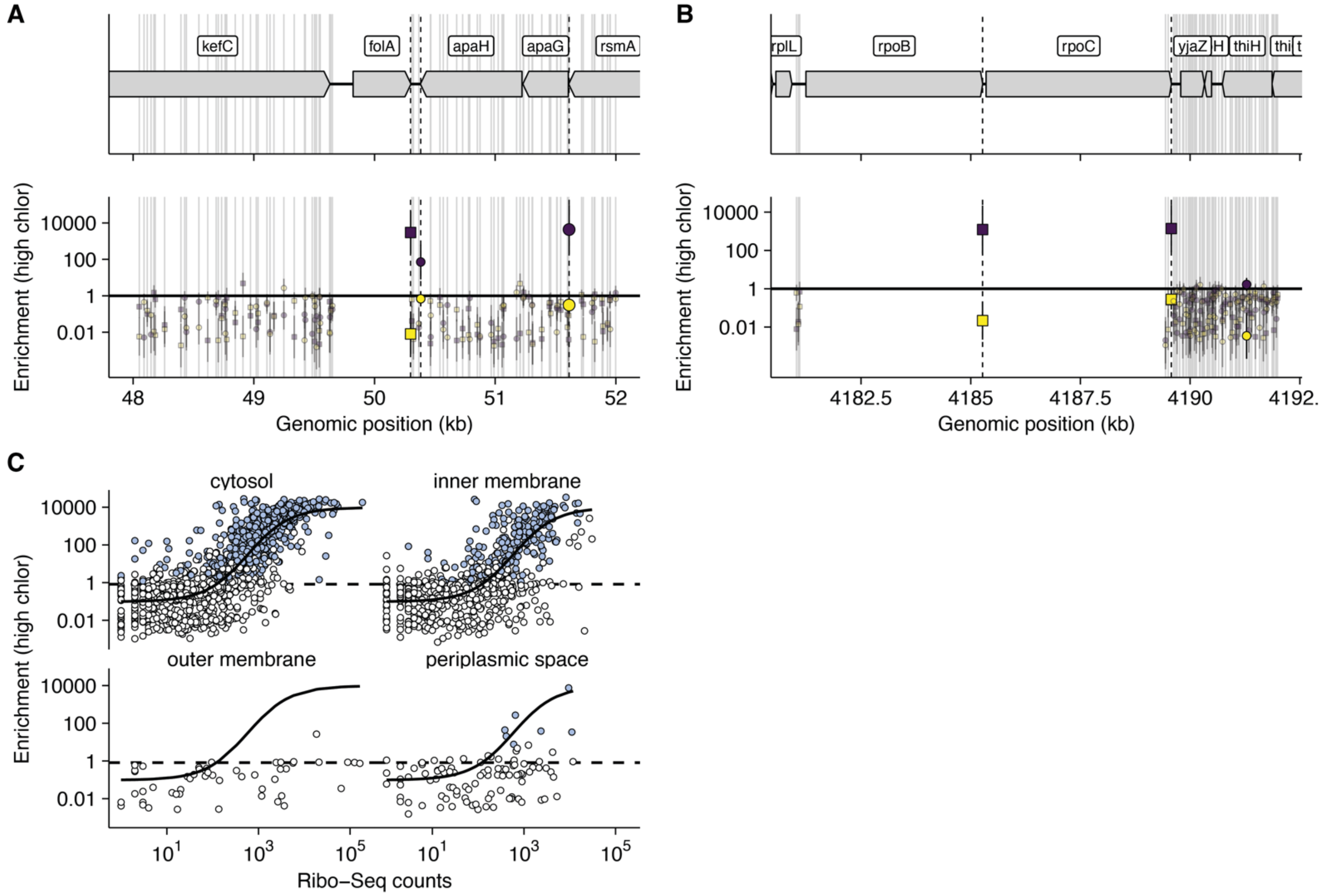


Fig. S4 Essential gene and localization constraints. A) An example essential gene is shown, folA. Symbols are same as main figure 2. B) Example essential genes, rpoB and rpoC are shown. C) Estimated enrichment vs. ribosome profiling data is shown for different localizations of annotated proteins. The top left panel is the same as main figure 2E, and the same cytosolic logistic model is applied to the other localizations to facilitate comparison. Blue points show significant (>75%) hits in high chloramphenicol.

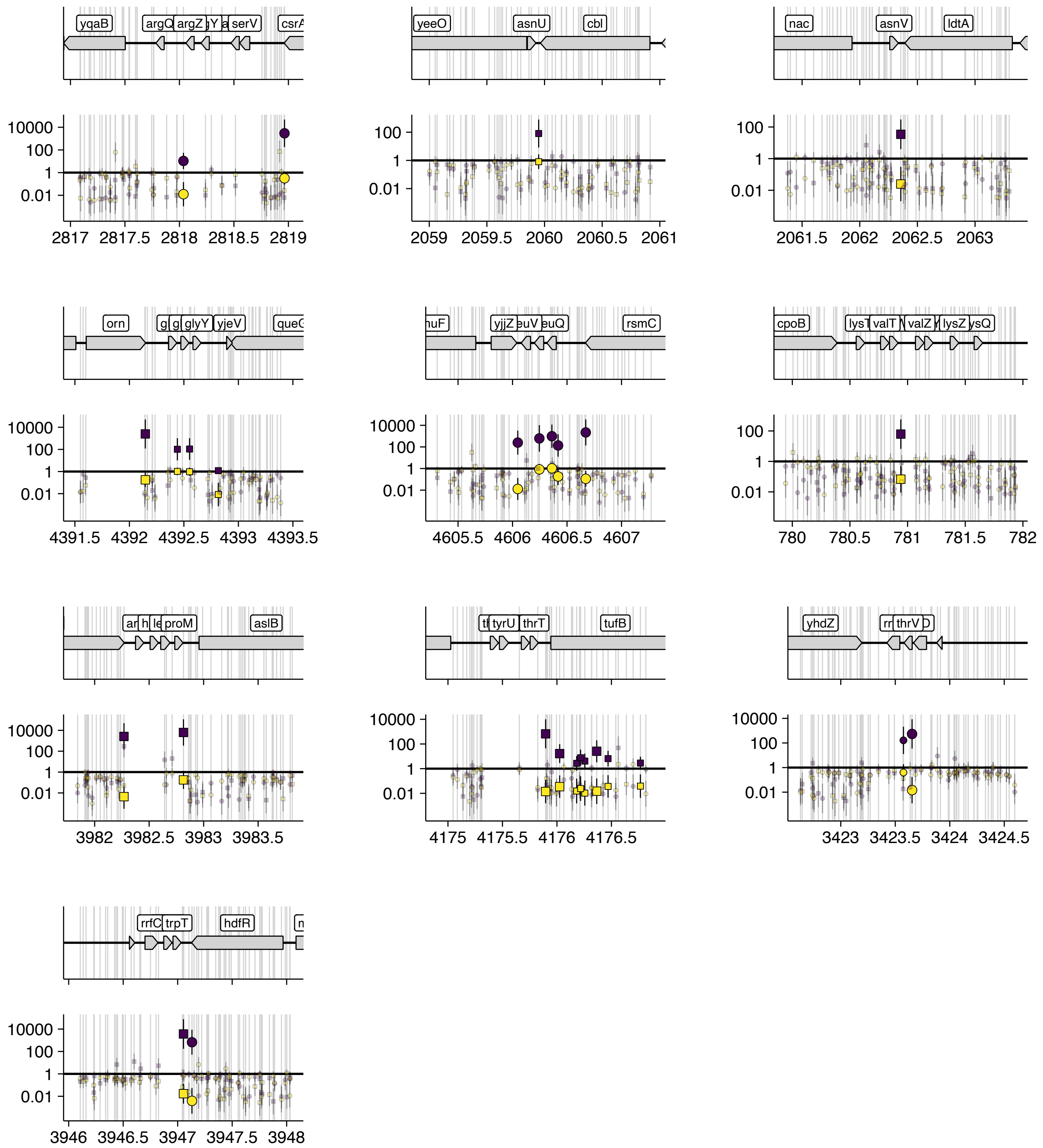


Fig. S5 Unannotated hits overlapping tRNA loci are shown. Symbols and coloring are the same as main figure 2.

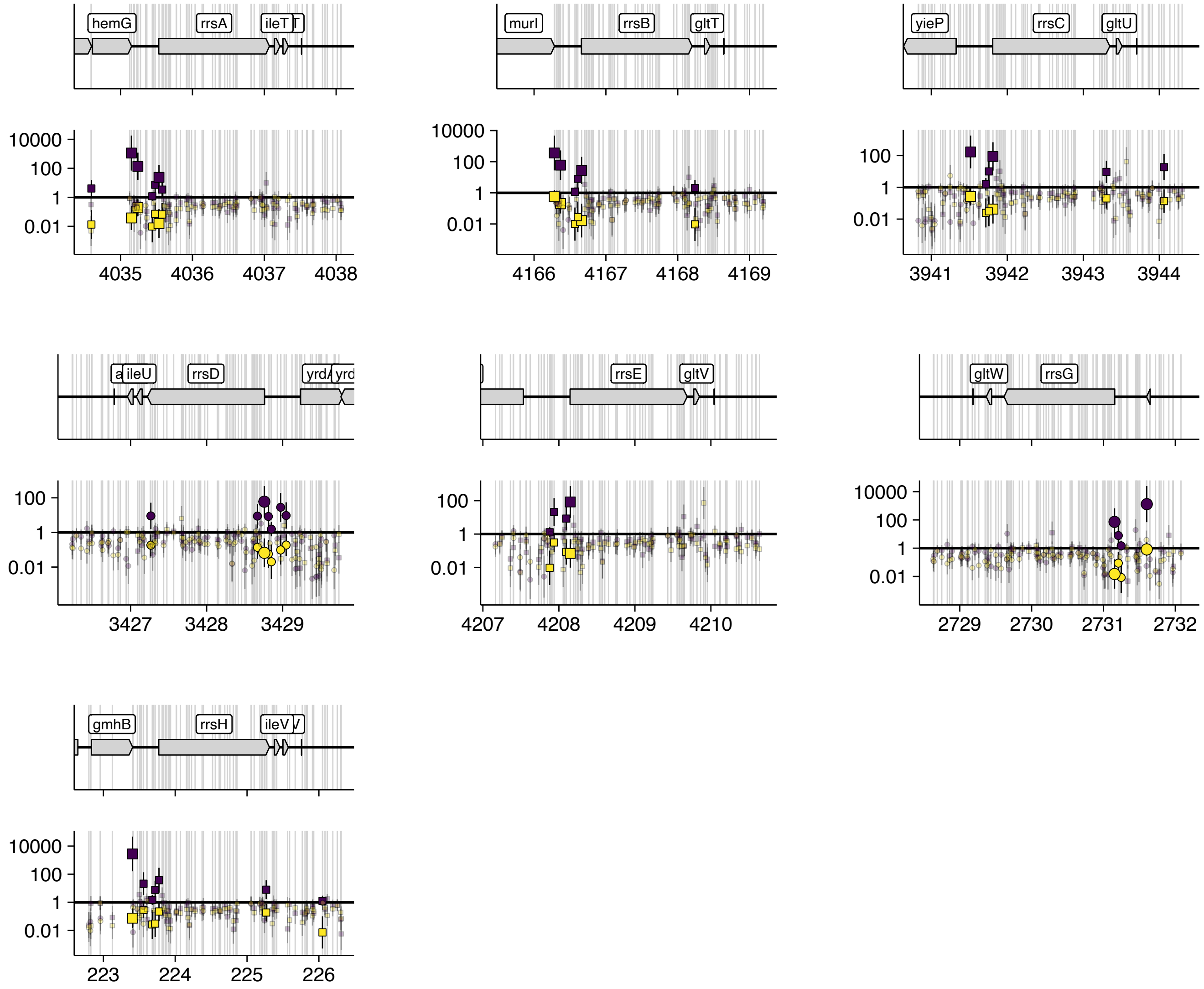


Fig. S6 Unannotated hits overlapping 16S rRNA loci are shown. Symbols and coloring are the same as main figure 2.

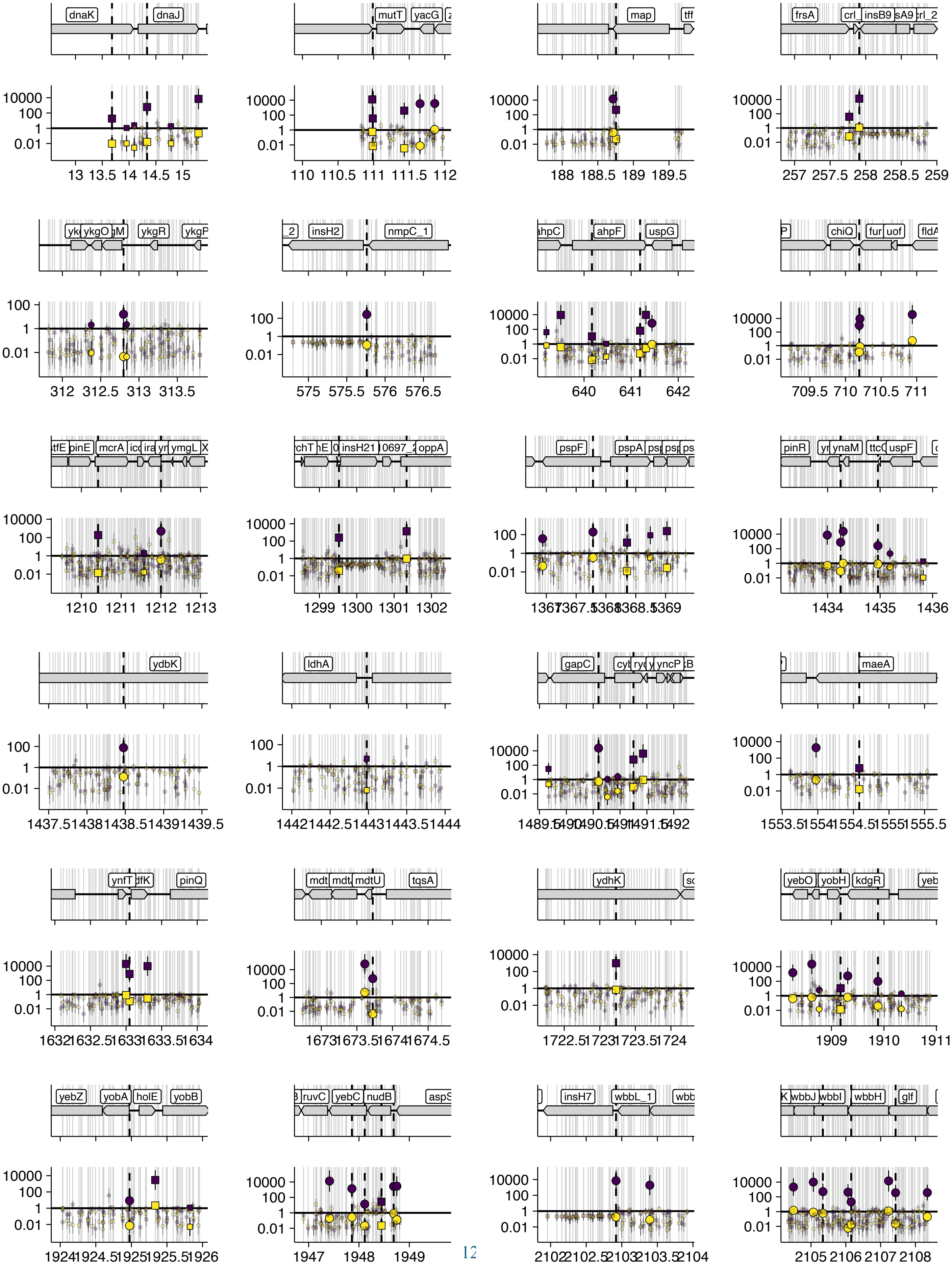
dnaK
dnaJ
mutT
yacG
map
tff
frsA
crl_
insB9
sA9
crl_2
ykgO
ykgR
ykgP
insH2
nmpC_1
ahpC
ahpF
uspG
chiQ
fur
uof
pinE
mcrA
ymgL
insH21
oppA
pspF
pspA
pinR
ynaM
uspF
ydbK
ldhA
gapC
yncP
maeA
ynfT
pinQ
mdtU
tqsA
ydhK
yebO
yobH
kdgR
yebZ
yobA
holE
yobB
ruvC
yebC
nudB
insH7
wbbL_1
wbbJ
wbbI
wbbH
glf

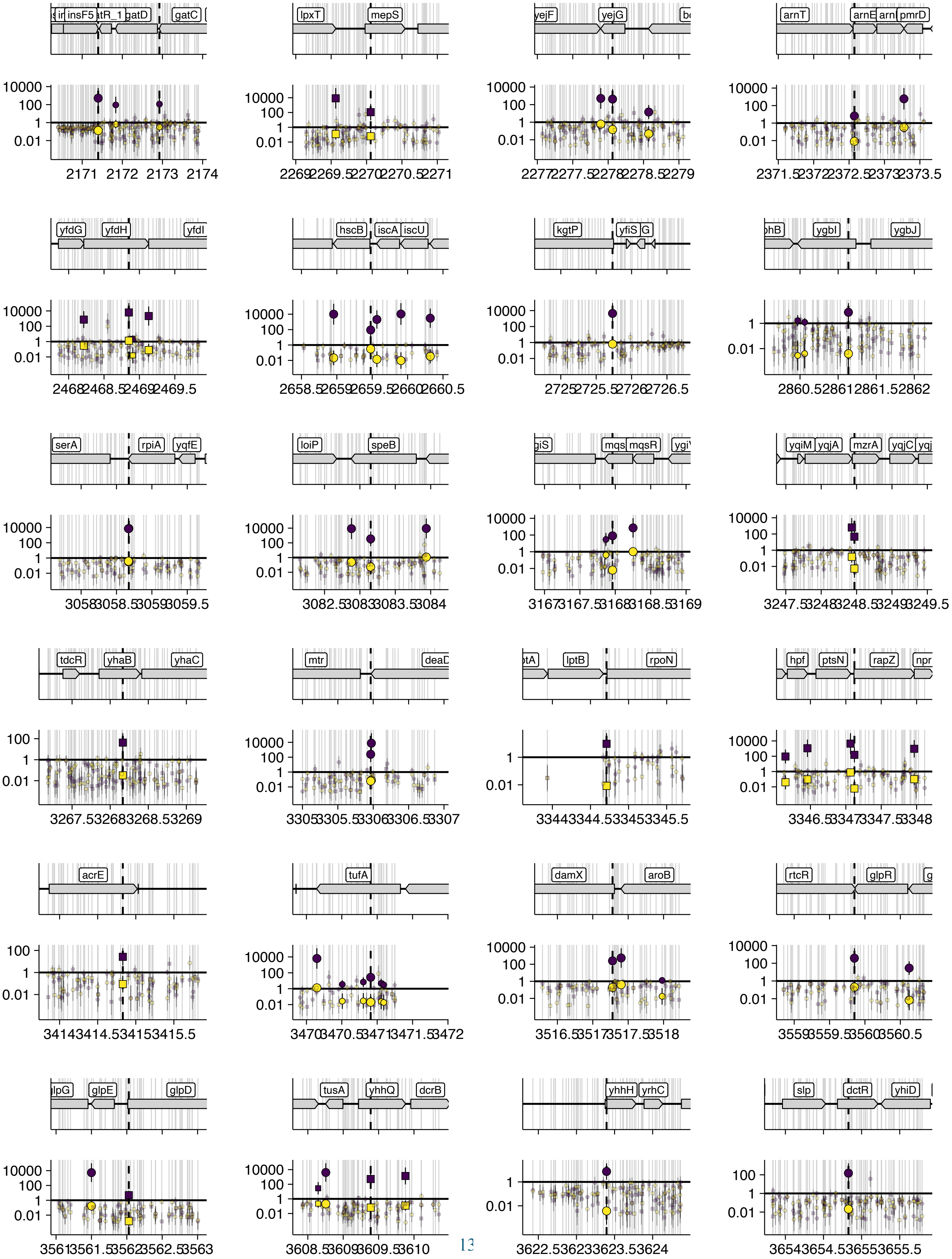
insF5
gatD
gatC
lpxT
mepS
yejF
yejG
arnT
pmrD
yfdG
yfdH
hscB
iscA
iscU
kgtP
yfiS
ygbI
ygbJ
serA
rpiA
yqfE
loiP
speB
mqsR
yqiM
yqjA
mzrA
yqjC
tdcR
yhaB
yhaC
mtr
lptB
rpoN
hpf
ptsN
rapZ
acrE
tufA
damX
aroB
rtcR
glpR
glpE
glpD
tusA
yhhQ
dcrB
yhhH
yrhC
slp
dctR
yhiD

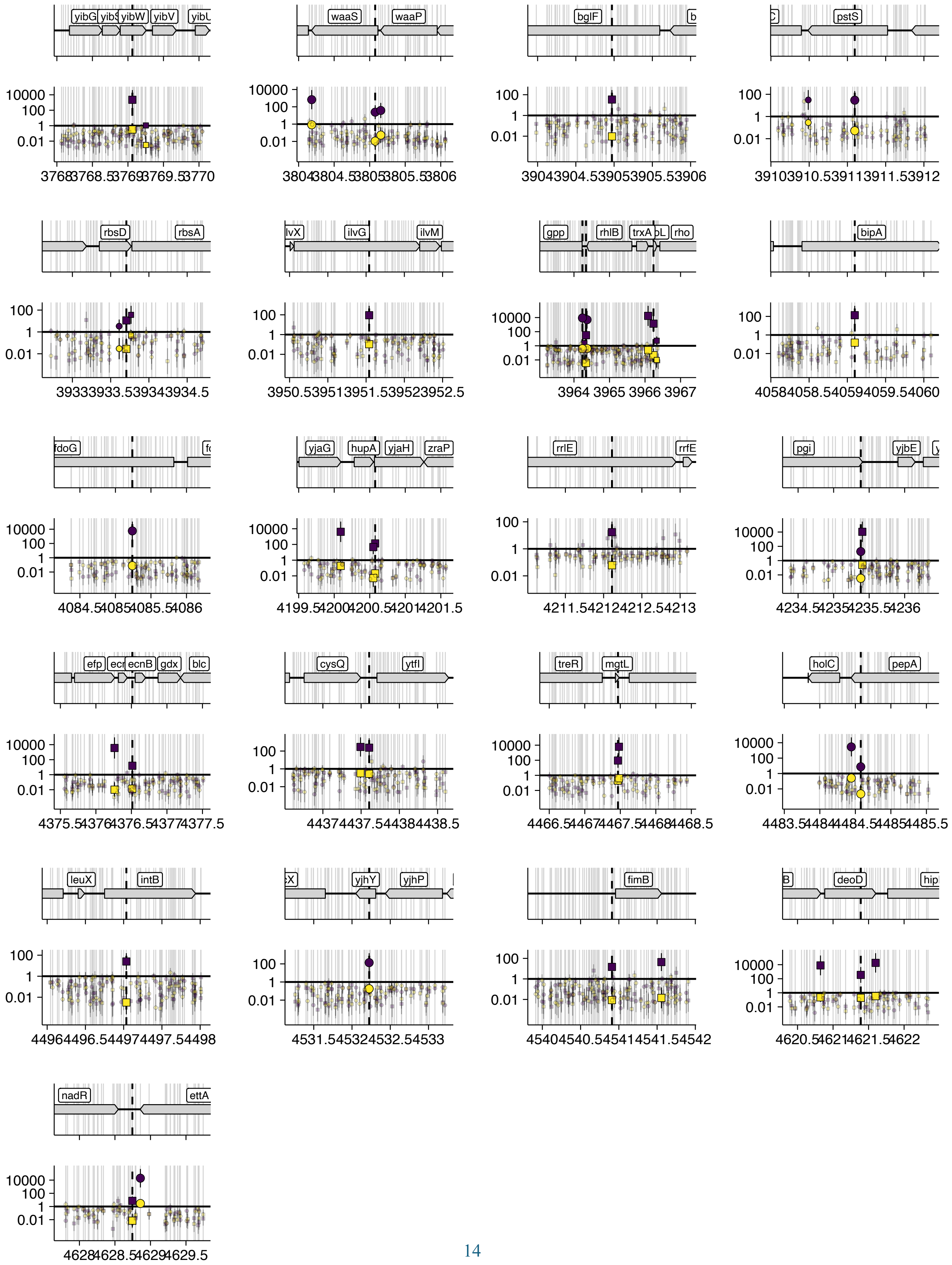
yibW
yibV
waaS
waaP
bglF
pstS
rbsD
rbsA
ilvG
ilvM
gpp
rhlB
trxA
rho
bipA
fdoG
yjaG
hupA
yjaH
zraP
rrlE
rrfE
pgi
yjbE
efp
ecnB
gdx
blc
cysQ
ytfI
treR
mgtL
holC
pepA
leuX
intB
yjhY
yjhP
fimB
deoD
nadR
ettA

Fig. S7 All other significant (>90% confidence) high chloramphenicol unannotated hits. Note that this includes several pseudogenes (e.g. ilvG) and a selenocysteine, fdoG. Coloring and symbols are the same as main figure 2.

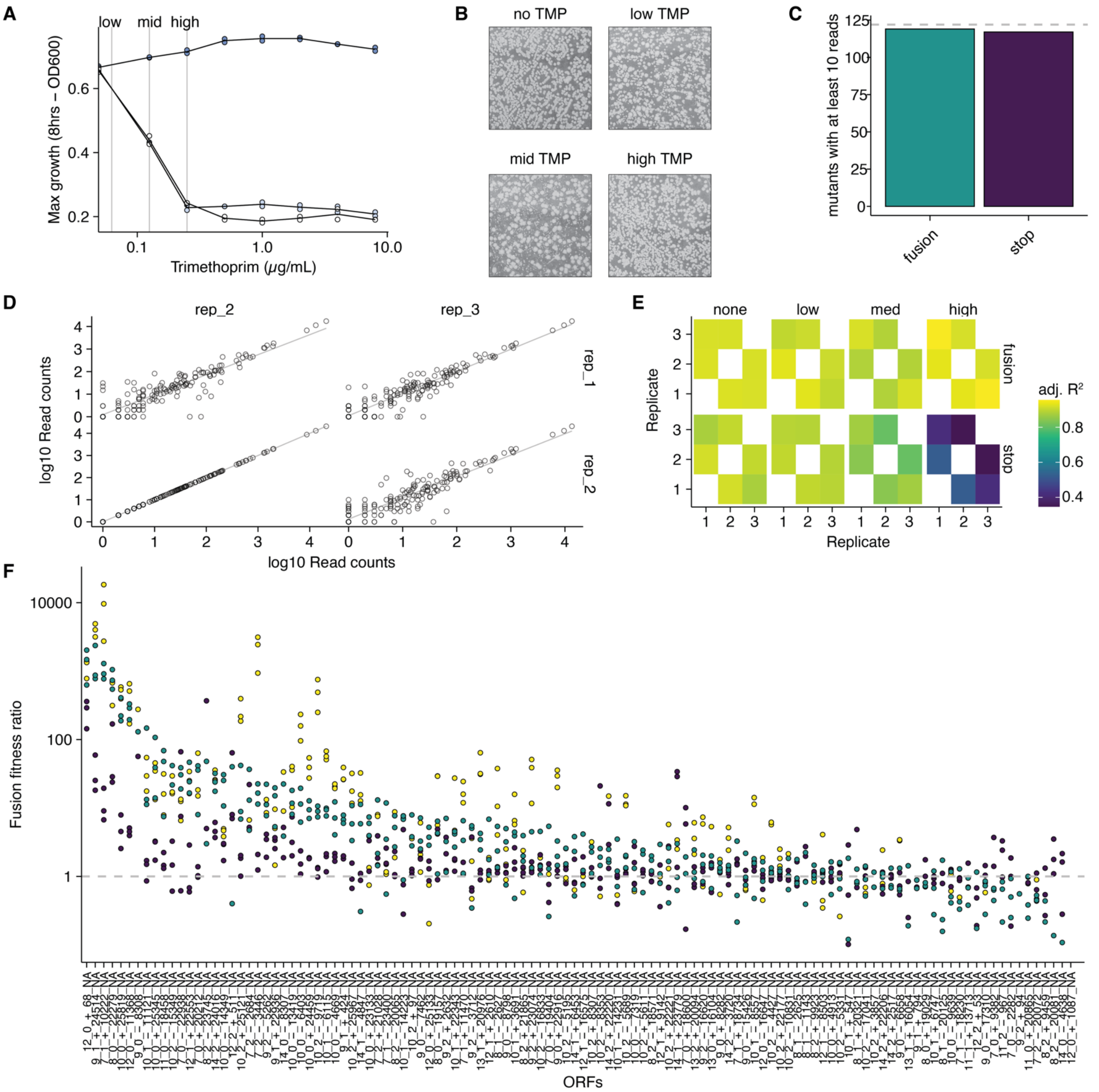


Fig. S8 mmDHFR fusion assay details. A) Max growth (OD600 for 8 hrs) is shown for acrZ-mmDHFR (blue) and acrZ-stop (white) at different concentrations of trimethoprim (TMP). Low, medium and high concentrations of TMP that are used throughout this work are shown as gray lines. B) Colony images on different TMP plates are shown with variable sized colonies. C) Mutants observed with at least 10 reads across the dataset are counted and compared to the 122 strain target (gray line). D) An example correlation for fusions in medium TMP is shown, with linear models displayed as gray lines. E) Adjusted $R^2$ values are shown for linear models fit to all replicate comparisons. F) FFR is shown with individual data points and ORF labels. Same as main fig 3C.

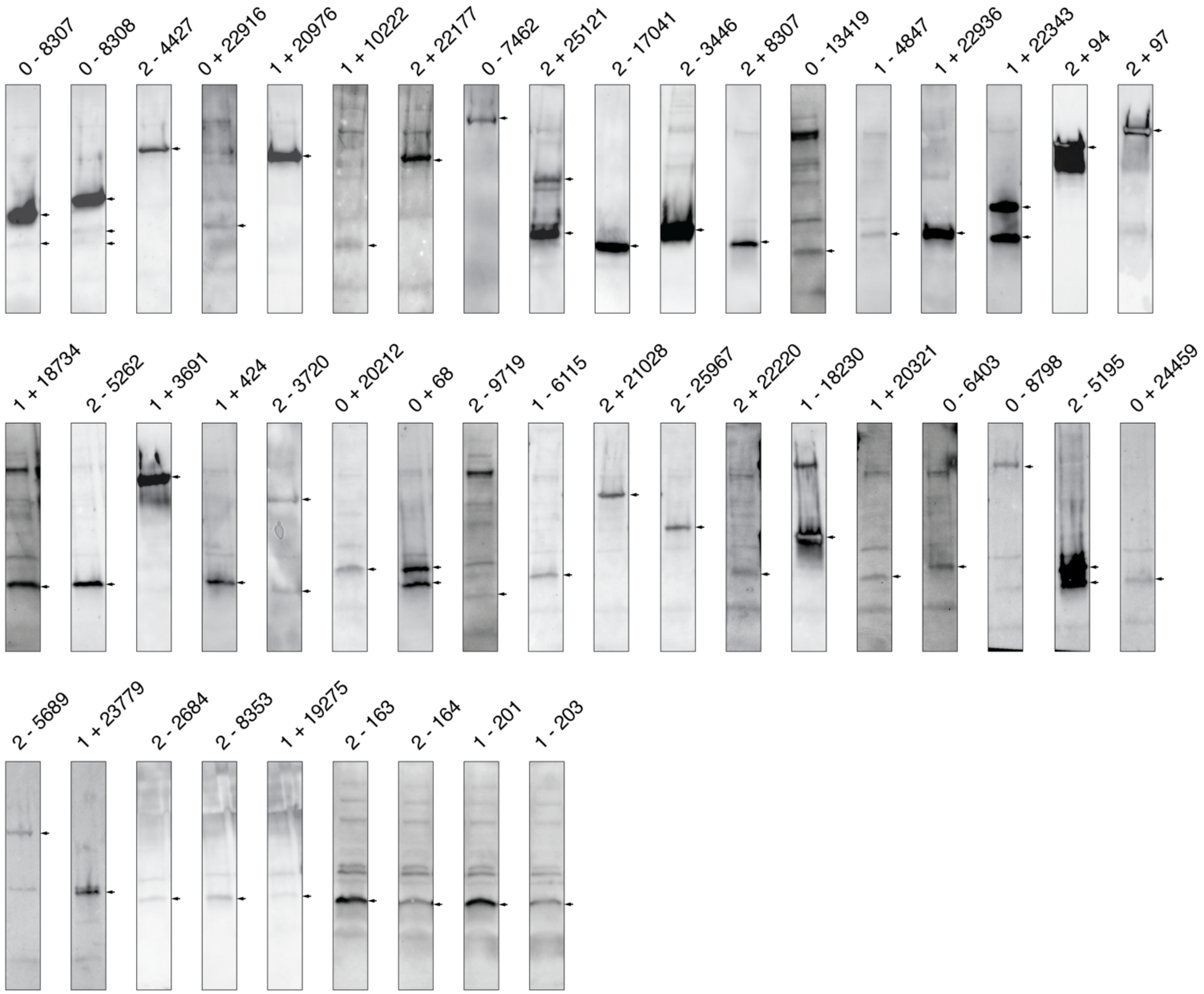


Fig. S9 Western blotted ORFs. A representative example of each observed western blot band is shown. Contrast has been adjusted to clearly show the observed band and each is highlighted with an arrow on the right side. Note that it is difficult to directly compare molecular weights across many different gels – raw western blot images are available at repo XYZ.

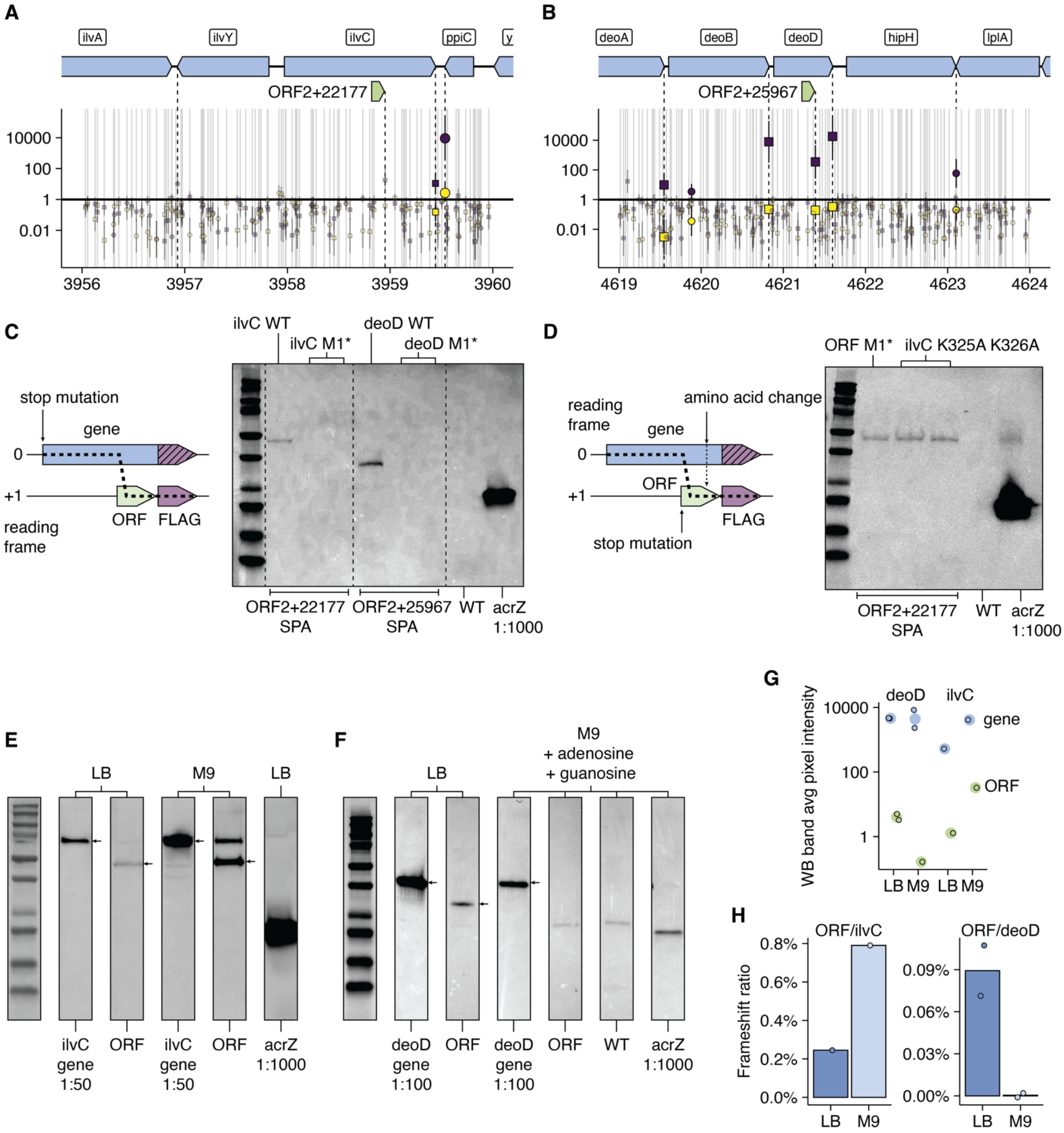


Fig S10. Ribosomal frameshift details. A-B) The chloramphenicol enrichment results are shown for the ilvC and deoD loci, with the sense overlapping ORF of interest highlighted. C) A western blot shows the putative frameshift product for ORFs overlapping ilvC and deoD and then shows the loss of the band upon inactivating the annotated gene start with a stop codon. D) A western blot shows that inactivating the ilvC overlapping ORF putative start with a stop codon does not eliminate the putative frameshift band. Point mutations changing the amino acid composition of ilvC (and the ORF) upstream of the SPA tag also do not eliminate the observed frameshift band. E-F) Western blots for ilvC, deoD and their frameshift products show different abundances in LB vs. M9 conditions. G) Semiquantitative western blot results (n=1 for ilvC and n=2 for deoD). H) The condition specific frameshifting ratio for ilvC and deoD in LB and M9 conditions.

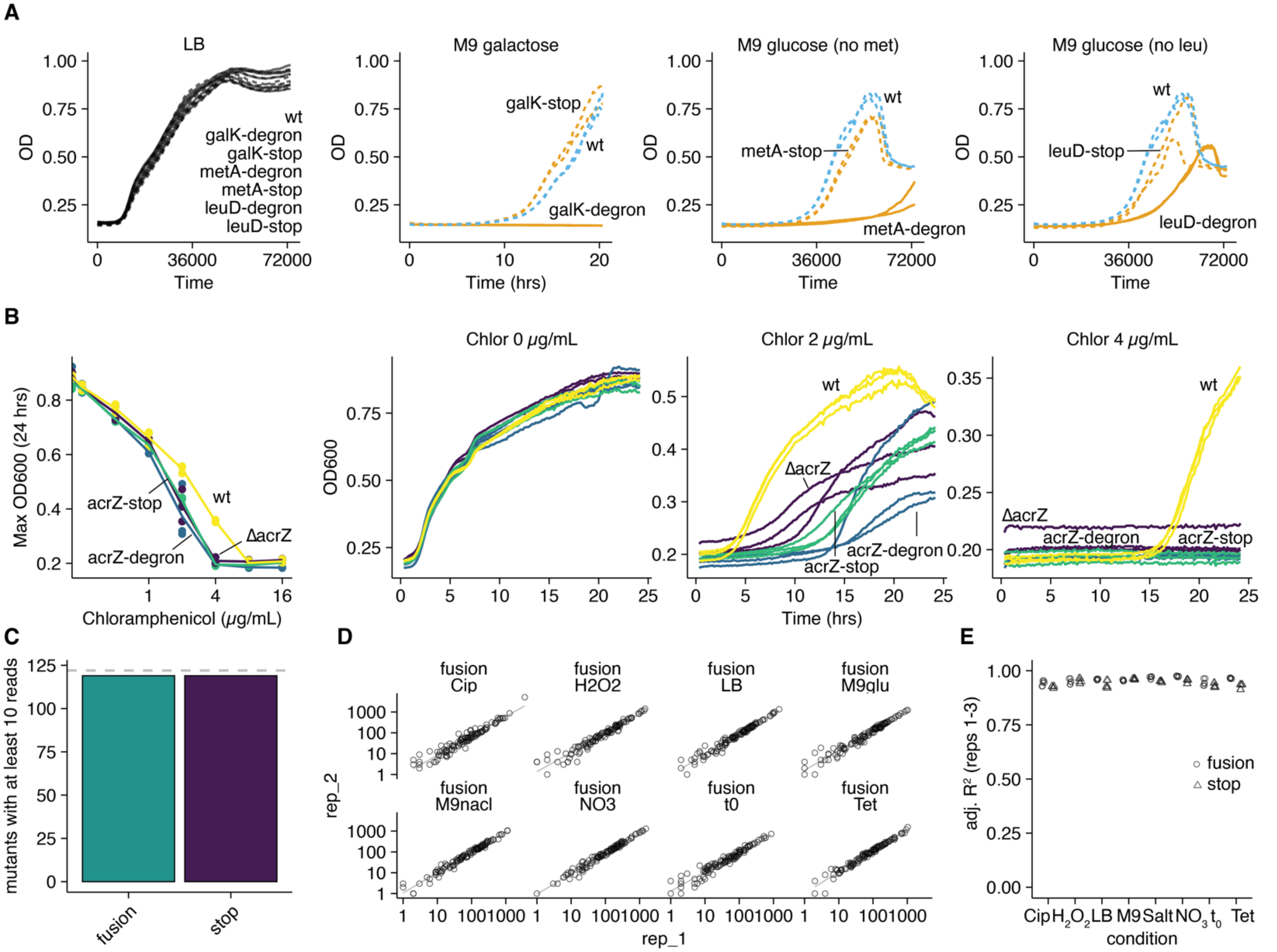


Fig. S11 Degron assay details. A) Growth curve of different degron and stop tagged strains growing in permissive medium, LB, and then selective media M9 galactose (galK), and M9 glucose (no added amino acids) for metA and leuD. B) Growth curves for acrZ deletion, degron and stop control strains. The left panel is a summary with the max OD600 during the growth curve and individual growth curves are shown to the right for increasing concentrations of chloramphenicol, which is natively affected by acrZ. C) Observed degron and stop control mutants (>10 reads) compared to the set of 122 targets (gray line). D) An example comparison of replicate degron data is shown for just two replicate fusions. Linear models are shown as gray lines. E) Linear models were fit to all replicate combinations and adjusted $R^2$ values are shown. Fusions are shown as open circles while stops are shown as open triangles.
